# The *In Vivo* Source of Type I and Type III IFNs is Pathogen Dependent

**DOI:** 10.1101/2021.10.05.463160

**Authors:** Marvin J. Sandoval, Hsiang-Chi Tseng, Heidi P. Risman, Sergey Smirnov, Qing Li, Jian-Da Lin, Amariliz Rivera, Russell K. Durbin, Sergei V. Kotenko, Joan E. Durbin

## Abstract

Type I (-α, β) and type III (-λ) interferons (IFNs) are produced in response to virus infection and upregulate a largely overlapping set of IFN stimulated genes which mediate the protective effects of these antiviral cytokines. *In vitro* studies have demonstrated the redundancy of these two cytokine families which activate the same transcription factor, IFN stimulated gene factor 3 (ISGF3), via distinct ligands and receptors. However, *in vivo*, these IFN types do have distinct functions based on receptor distribution, but also ligand availability. Using a newly generated IFN-λ reporter mouse strain we have observed that both type I and type III IFNs are produced in response to respiratory tract infection by Newcastle disease virus (NDV) and influenza A virus (IAV). In the case of NDV these IFNs are synthesized by different cell types. Type I IFNs are produced primarily by alveolar macrophages, type III IFNs are made only by epithelial cells, and production of either is dependent on MAVS. While epithelial cells of the respiratory tract represent the primary target of IAV infection, we found that they did not significantly contribute to IFN-λ production, and IFN-λ protein levels were largely unaffected in the absence of MAVS. Instead we found that pDCs, a cell type known for robust IFN-α production via TLR/MyD88 signaling, were the major producers of IFN-λ during IAV infection, with pDC depletion during influenza infection resulting in significantly reduced levels of both IFN-α and IFN-λ. In addition, we were able to demonstrate that pDCs rely on type I IFN for optimal IFN-λ production. These studies therefore demonstrate that the *in vivo* producers of Type III IFNs in response to respiratory virus infection are pathogen dependent, a finding which may explain the varying levels of cytokine production induced by different viral pathogens.

## Introduction

The antiviral activity of the type I interferons (IFN-α/β) has been appreciated since their discovery in 1957 [1, 2]. The more recent discovery of type III IFNs (IFN-λ) [3] [4], a new IFN family with its own distinct receptor but mediating the same antiviral response, has led to a reevaluation of our understanding of innate immune responses. The IFN-λ receptor, comprised of the IFN-lambda receptor (IFNLR) chain and the IL-10R2 chain, is expressed on most epithelial cells, but not ubiquitously found on all cell types like the IFN-alpha receptors (IFNAR) 1 and 2. The type III IFNs act to protect mucosal surfaces, the point of entry for many viral pathogens, but their source(s), and the stimuli leading to their production are not yet well characterized [7, 8]. To better understand the relative contributions of type I and type III IFNs to antiviral protection, we generated an IFN-λ reporter mouse, *Ifnl2 ^gfp/gfp^*, and used this strain to examine the relative contribution of various cell types to the production of these cytokines. This approach was inspired by the work of Kumagai *et al*. [5] who demonstrated that alveolar macrophages produce the bulk of IFN-α following pulmonary infection with Newcastle disease virus (NDV) using mice expressing GFP under the control of the *Ifna6* promoter. Similar results were obtained following infection of these *Ifna6^gfp/+^* mice with respiratory syncytial virus (RSV) [6].

Multiple studies have demonstrated that type I and type III IFNs are induced *in vitro* by the same stimuli, e.g. virus infection and TLR ligand exposure [7] [8], but it is not yet known if induction consistently follows the same pathways following virus infection *in vivo*. Making use of *Ifnl2^gfp/gfp^* mice we have confirmed Kumagai’s observation that alveolar macrophages are the major source of IFN-α during NDV infection, but also observed that this cell type produces no IFN-λ. IFN-λ in this infection is entirely derived from respiratory epithelial cells in a MAVS-dependent manner. The finding that IFN-λ is preferentially produced by polarized epithelial cells *in vitro* [9] has also been observed in mouse models [10]. These published results support the notion that, despite their activation of the same antiviral pathway, IFN-α/β and IFN-λ are produced by different cell types in response to the same stimulus.

In this study we used the *Ifnl2^gfp/gfp^* reporter mice to test the generality of this hypothesis with a number of viruses. In both the NDV and rhesus rotavirus (RRV) infections we observed IFN-λ production by the IFN-λ responsive cells lining the target tissue, emphasizing the important role for this cytokine in protection of the epithelial barrier. This pattern of IFN expression was not recapitulated in the murine model of influenza A virus (IAV) infection, where a significant proportion of IFN-λ induced by infection was produced by pDCs in a MAVS-independent manner. Taken together our studies show that the source of type I and type III IFNs *in vivo* is pathogen dependent, and not defined by the route of infection or the tropism of the infecting virus.

## Results

### Generation and Characterization of the IFN-λ reporter mouse

To identify IFN-λ-producing cell populations *in vivo*, we generated an IFN-λ reporter mouse by replacing the coding region of *ifnl2* with an *eGFP-Neo* cassette through homologous recombination, while maintaining the *ifnl2* promoter and UTR regions intact (Fig 1A). The FRT sites flanking the neo gene allowed its removal by crossing mice carrying the targeted allele with mice expressing the FLP-recombinase to generate animals lacking the Neo gene while simultaneously carrying the GFP-targeted allele. Mice heterozygous for the eGFP-targeted allele were crossed to generate mice homozygous for the targeted allele. Homozygous *ifnl2*^eGFP/eGFP^ knockin mice can no longer produce IFN-λ2, but still retain functional *ifnl3* gene loci. Genotypes were determined by conventional PCR using primers targeting the IFN-λ locus and the eGFP transgene (Fig 1A, B).

**Figure 1.**
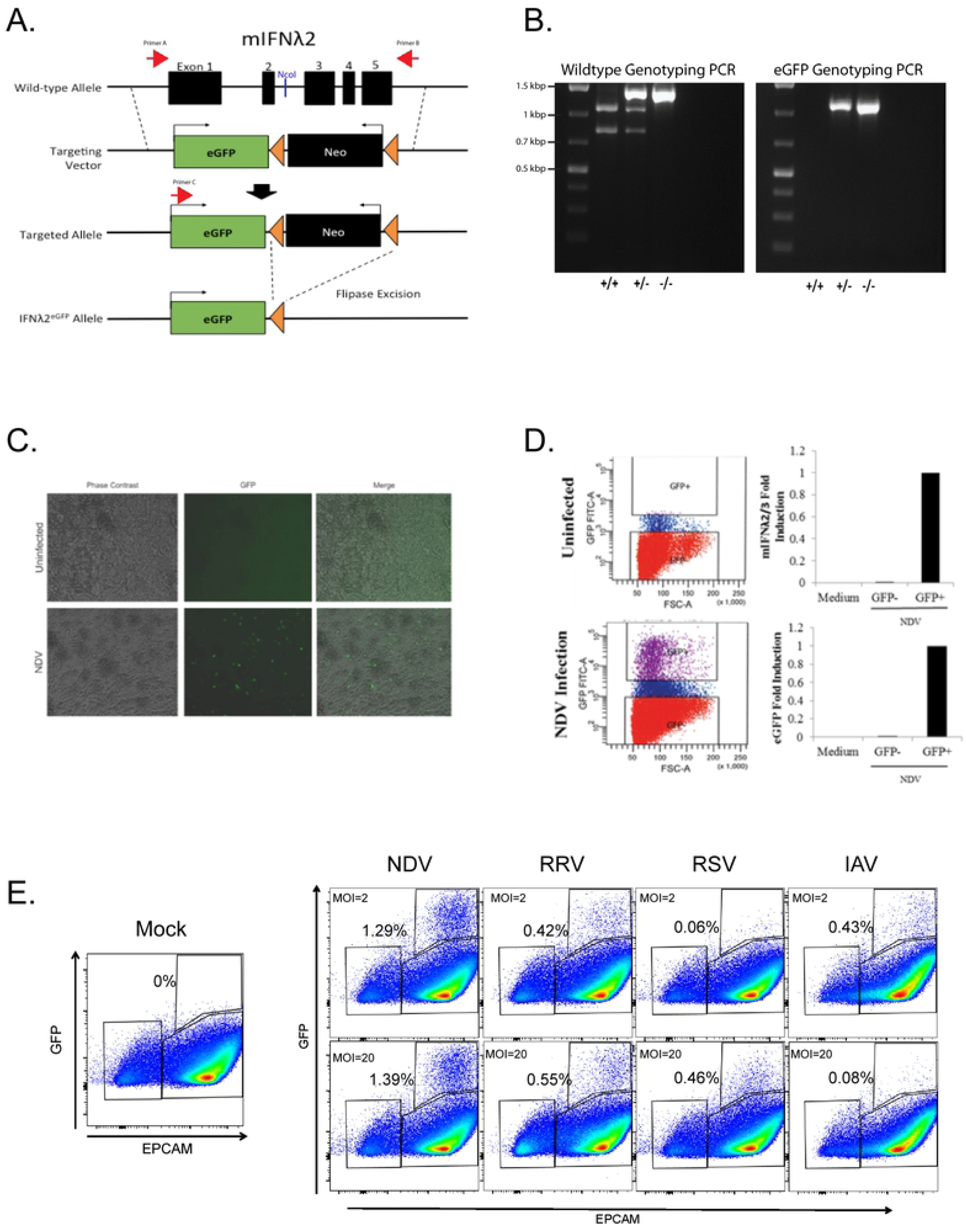
Generating the IFN-λ Reporter Mouse. (A) The targeting vector carrying *eGFP* and *Neo* genes was inserted into the *Ifnl2* locus by homologous recombination, resulting in complete replacement of the *Ifnl2* coding region with the *eGfp* and *Neo* genes. Excision of the *Neo* gene was carried out by crossing mice carrying the targeted allele with Flipase mice, resulting in deletion of the *Neo* gene and generation of the *Ifnl2^+/gfp^* allele. (B) PCR products generated with primers shown in (A), and digested with Nco1, yielded unique banding patterns for tail DNA obtained from WT 129 SvEv mice (+/+), heterozygous *Ifnl2^+/gfp^* (+/-), and homozygous *Ifnl2^gfp/gfp^* IFN-λ reporter animals. (C) Pictured are phase contrast and GFP immunofluorescence images (200x) of uninfected or NDV-infected, FLT3L cultured, bone marrow derived DCs from *Ifnl2^gfp/gfp^* mice. Images are representative of three independent experiments. (D) NDV-infected FLT3L cultured, bone marrow derived DCs were cell sorted into GFP+ and GFP- fractions that were then assayed by qRT-PCR for the presence of GFP and IFN-λ3 transcripts. Data are representative of two independent experiments. (E) Flow cytometry dot plots show GFP expression in cultures of primary kidney epithelial cells (gated as EPCAM+ cells) derived from *Ifnl2^gfp/gfp^* mice that were mock-infected or infected with Newcastle disease virus (NDV), rhesus rotavirus (RRV), respiratory syncytial virus (RSV), or influenza A virus (IAV) at the indicated MOIs for 24 hrs. Data are representative of two independent experiments.

Human pDCs secrete high levels of both type I and type III IFNs when stimulated by TLR ligands or virus infection [11], so characterization of the reporter strain was initiated by Newcastle disease virus (NDV) infection of Flt3-ligand (FLT3L) cultured bone marrow derived dendritic cells. NDV is an avian paramyxovirus known to induce high levels of type I IFNs in many cell types including murine pDCs [5, 12], so we used this system to ask whether GFP expression correlated with IFN-λ production upon induction by virus infection.

GFP expression following virus inoculation was detected by fluorescence microscopy (Fig 1C), as well as flow cytometry (Fig 1D). A Fluorescence Activated Cell Sorter (FACS) was used to separate NDV-treated, FLT3L-cultured DCs into GFP- and GFP+ populations for RNA extraction and subsequent qRT-PCR analysis of IFN-λ3 and eGFP transcripts (Fig 1D). This analysis confirmed that IFN-λ3 and eGFP transcripts were enriched only in GFP+ cells, demonstrating that GFP fluorescence from reporter cells reflected IFN-λ production.

We also explored GFP induction following virus infection of epithelial cells. Primary kidney epithelial cells derived from *ifnl2^gfp/gfp^* reporter mice were cultured *in vitro* and infected with NDV, respiratory syncytial virus (RSV), rhesus rotavirus (RRV) or influenza A virus (Fig 1E). At 24 hours post-infection, all infected cultures contained GFP+ cells, but for all viruses the percentage of GFP+ cells was relatively small, even at higher MOIs. A similar effect was observed in FLT3L cultured DCs (data not shown), suggesting that during a virus infection, not all cells capable of producing IFN-λ go on to do so as has been found for IFN-α [13].

### A small population of CD8α DCs produce IFN-λ in response to poly I:C treatment *in vivo*

Production of IFN-λ by different cell types has been detected in response to a wide variety of stimuli or pathogens *in vitro* [14–16]. Taken together, these publications demonstrate that many cell types have the capacity to produce type III IFNs under the appropriate conditions. However, reports investigating the cellular sources of IFN-λ in a whole animal in response to TLR ligands or virus infections are few. One such study identified splenic CD8α+ dendritic cells as the main producers of IFN-λ in mice that were treated intravenously (i.v.) with the TLR3 agonist, poly I:C [17]. We carried out a similar experiment using our IFN-λ reporter mouse model to determine whether we could confirm that result by following GFP expression. Cohorts of *Ifnl2^gfp/gfp^* reporter mice were injected i.v. with 100 μg of poly I:C or PBS, and their splenocytes harvested six hours later. Using antibodies to the cell surface markers CD11b, CD11c, CD8α, B220, and Siglec H, we identified pDC, CD8α DC, and cDC subsets, and found, as expected, that GFP expression was mainly confined to the CD8α DC populations (Fig 2). Interestingly, IFN-λ-producing (GFP+) CD8α DCs made up only 6% of the entire CD8α DC population; GFP expression by pDCs and cDCs was minimal. This result is consistent with the observation of Lauterbach *et al*. [17] that CD8α DCs are by the far the most significant contributor to IFN-λ production in response to poly I:C, supporting the use of the *Ifnl2^gfp/gfp^* mouse strain to determine the source of IFN-λ *in vivo*. No upregulation of GFP was found in animals receiving PBS.

**Figure 2.**
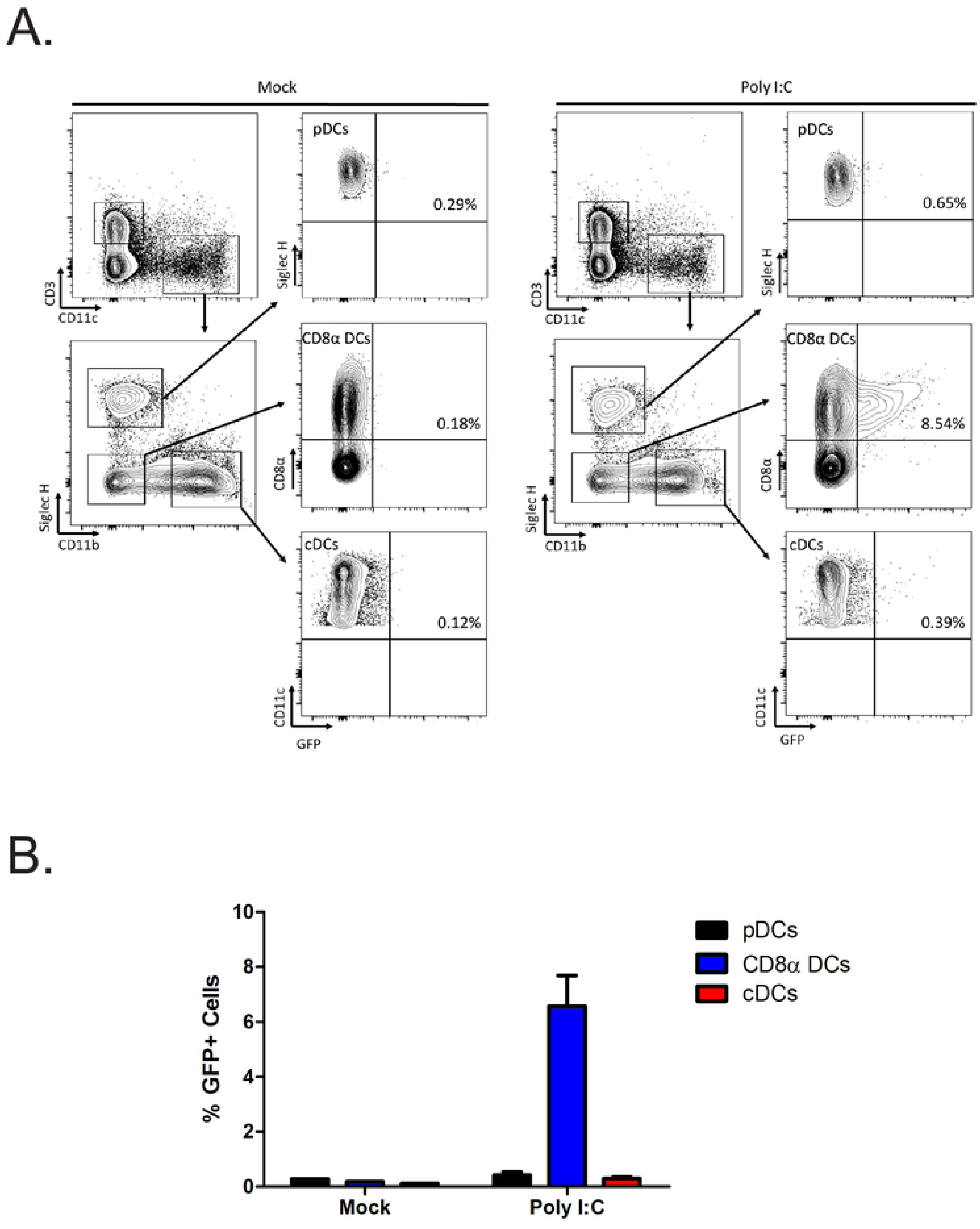
CD8α DCs are the predominant IFN-λ-producing cells in spleens of mice treated intravenously with poly I:C. (A) Flow cytometry analysis of splenic populations from IFN-λ reporter mice mock-treated (left) or i.v. treated with poly I:C (right) for 6 hrs. Cells were gated on CD11c while simultaneously excluding CD3+ cells to exclude T cells. Dendritic cell populations were then identified as follows: pDCs (CD11c+/Siglec-H+/CD11b-); CD8α DCs (Siglec-H-/CD11b-/CD8α+); cDCs (Siglec-H-/CD11b+/CD11c+). (B) Mean percentages, with SEM, of GFP+ cells in populations shown in (A). Data are representative of two independent experiments.

### Intestinal epithelial cells produce IFN-λ in response to rotavirus infection *in vivo*

As both type I and type III IFNs are produced in response to many different virus infections *in vivo* [14] [7], we wished to investigate the cellular sources of type III IFNs during virus infection of mucosal surfaces such as the gastrointestinal tract and the respiratory tract. Rotavirus is a double-stranded RNA virus that primarily infects intestinal epithelial cells and is one of the leading causes of severe diarrhea in young children in the developing world [18]. Several studies have shown the importance of IFN-λ responses, along with type I IFNs, in controlling rotavirus replication [19, 20]. Here, we explored the *in vivo* source of IFN-λ during rhesus rotavirus (RRV) infection of suckling mice. The kinetics of IFN induction by RRV was first determined using Mx2-Luciferase mice, a reporter mouse strain that allows visualization and quantification of IFN responses by measuring luciferase activity driven by the promoter of the interferon-stimulated gene Mx2 [21]. By this approach, a robust interferon response was detected as early as 12 hours post-infection, peaking at 24 hours, and subsiding after 48 hours (Fig 3A&B). Consistent with this observation, IFN-λ protein was detected in the small intestine of rotavirus-infected WT 129SvEv mice 24 hours post-infection by ELISA (Fig 3C). To determine the source of IFN-λ induced by RRV, intestines were harvested from infected IFN-λ reporter mice 24 hours post-inoculation and analyzed by immunohistochemistry. Formalin-fixed, paraffin-embedded tissue sections from mock-infected or RRV-infected IFN-λ reporter mice were stained with antibodies against GFP and RRV antigens (Fig 3E). GFP+ cells were present only in tissues harvested from RRV-infected IFN-λ reporter mice, and absent from the intestines of mock-infected IFN-λ reporter mice, or RRV-infected WT 129SvEv animals. GFP expression in RRV-infected IFN-λ reporter mice was found exclusively in the intestinal epithelial cells. RRV-infected epithelial cells were found in both RRV-infected IFN-λ reporter mice and RRV-infected WT controls, confirming successful infection of these animals. These results support the hypothesis that epithelial cells are the primary source of IFN-λ during RRV infection. As IFN-λ reporter mice lack the *ifnl2* locus but have an intact *ifnl3* locus, we compared RRV titers from the small intestine of IFN-λ reporter and WT 129SvEv animals, and found no difference in the amount of virus detected (Fig 3D). Thus the reporter mice are phenotypically normal, without enhanced susceptibility to virus infection or elevated virus loads.

**Figure 3.**
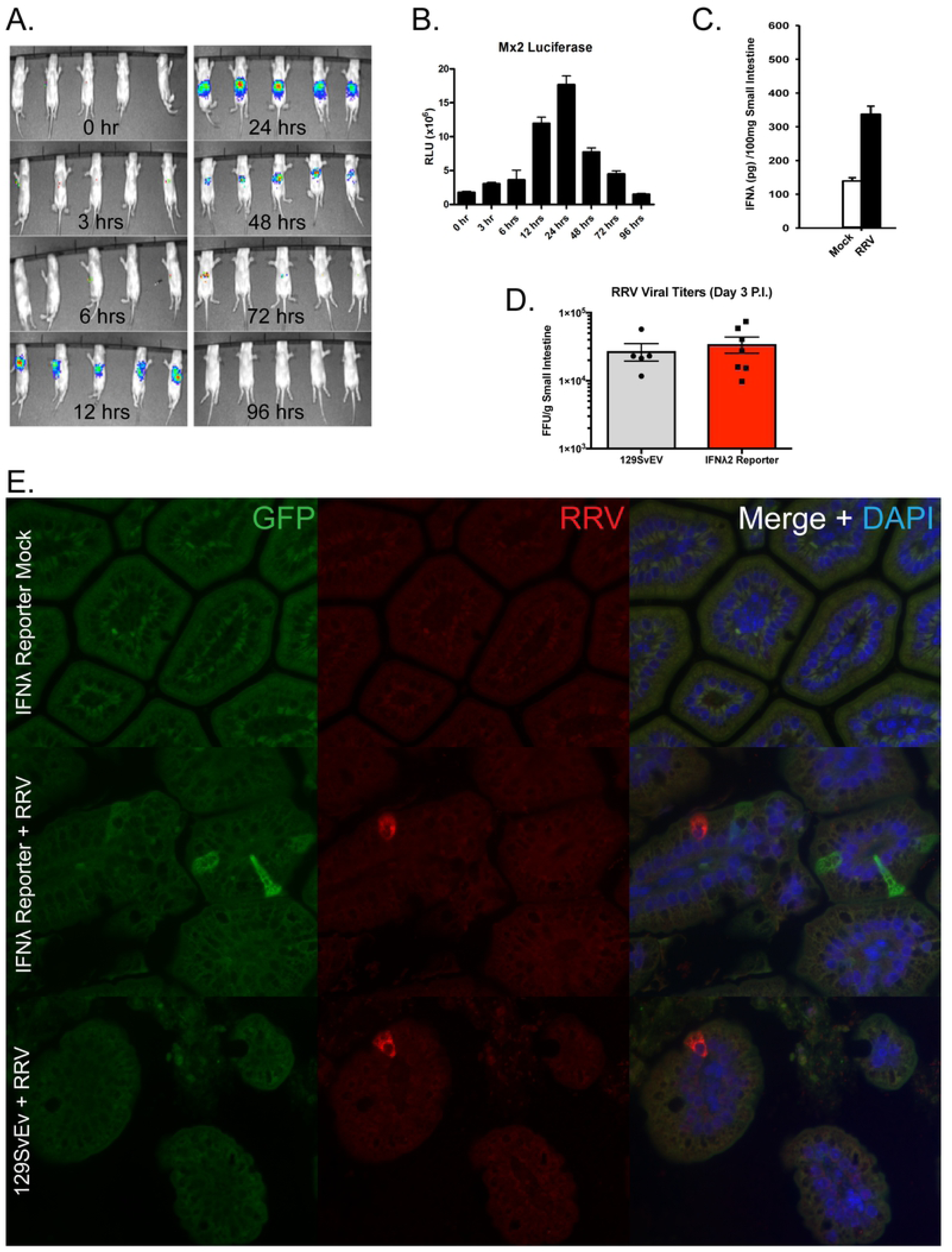
Epithelial cells are the predominant IFN-λ-producing cell population in the small intestine in response to RRV. (A) Interferon responses of 6-day old, RRV-infected Mx2-Luciferase transgenic mice were quantitated by IVIS at the indicated timepoints. (B) Graphical representation of mean luciferase activity values and SEM from two independent experiments. (C) ELISA measurements of IFN-λ protein in homogenates of intestinal tissue harvested from 6- day old WT C57BL/6 pups, 24 hours after RRV infection. (D) Comparison of RRV viral titers, measured by fluorescent-focus assay, in the small intestine of 6-day old WT 129 SvEV mice and IFN-λ reporter mice 3 days post-RRV infection. Data are pooled from two independent experiments. (E) Images of formalin-fixed paraffin-embedded (FFPE) small intestinal tissue sections from mock- or RRV-infected IFN-λ reporter pups, and RRV-infected WT pups, were captured 24 hours post-infection. Sections were probed with anti-GFP antibodies (green), anti-RRV sera (red) and DAPI (blue) and visualized by fluorescence microscopy. Images are representative of three independent experiments.

### Airway epithelial cells produce IFN-λ in response to NDV infection *in vivo*

Next, we sought to investigate IFN-λ production during virus infection of the respiratory tract, another mucosal compartment that serves as a major portal for virus entry. Our lab and others have reported that both type I and type III IFNs are produced in response to respiratory virus infections [22, 23]. In the lung, as in the intestine, both IFN types appear to be important for antiviral protection of the respiratory tract, as mice deficient in both type I and type III IFN receptors demonstrate significantly elevated viral burdens upon virus challenge [24, 25]. To determine whether the airway lining cells are major IFN-λ producers during respiratory virus infection *in vivo*, we inoculated IFN-λ reporter mice with Newcastle disease virus (NDV), a virus known to induce high levels of type I IFNs in mammals [26, 27]. At 24 hours post-infection, lungs from mock-infected or NDV-infected mice were harvested and processed to generate single cell suspensions. IFN-λ-producing, epithelial cell adhesion molecule (EpCAM)+ epithelial cells were assayed for GFP expression by flow cytometry. Only epithelial cells from NDV-infected reporter mice were GFP+ (Fig 4A), with approximately 15% of lung epithelial cells in the IFN-λ reporter mice expressing IFN-λ in response to infection *in vivo* (Fig. 4B). ELISA measurements of IFN-λ protein present in bronchoalveolar lavage samples (BALs) from these same animals also demonstrated robust IFN-λ secretion in response to NDV infection, with > 10 ng/ml of IFN-λ protein detected (Fig 4C).

**Figure 4.**
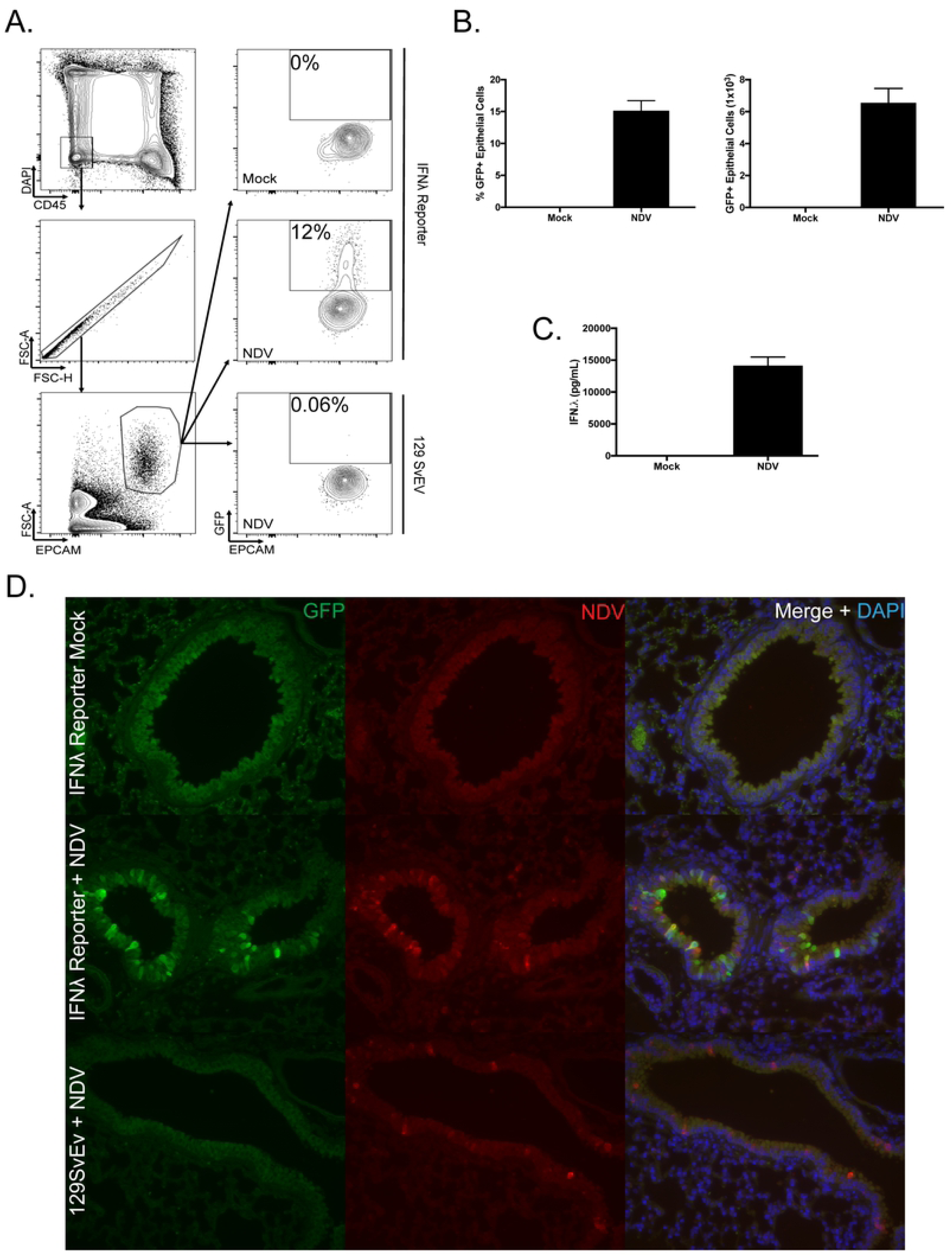
Epithelial cells are the predominant IFN-λ producers in response to NDV infection in vivo. (A) GFP expression was assayed by flow cytometry of cells obtained from collagenase-digested lungs of IFN-λ reporter mice which were mock-infected or infected intranasally with NDV for 24 hrs. Epithelial cells were defined as CD45-, EPCAM+. WT 129SvEV mice were included as controls for GFP expression, gating in the same manner. (B) Graphical representation of mean percentage of GFP+ epithelial cells from each treatment cohort. Error bars represent SEM. (C) IFN-λ protein levels in BALs harvested at 24 hours from mock-infected or NDV-infected reporter mice were determined by ELISA. (D) Lung sections from uninfected IFN-λ reporter mice, and NDV-infected WT and NDV-infected IFN-λ reporter mice were stained with anti-GFP antibodies (green), anti-NDV antibodies (red), and DAPI (blue). Data are representative of three independent experiments.

As with our rotavirus studies, we looked for GFP+ cells in tissue sections prepared from IFN-λ reporter mice that were mock or NDV-infected. As before, lung tissue sections were stained using antibodies against GFP and viral antigens (Fig 4D). This histological analysis confirmed our flow cytometry results, showing that GFP expression was limited to the columnar epithelial cells that line the bronchi and bronchioles of the lung. The same cell type expressed both NDV and GFP protein in infected lungs. Interestingly, while rare cells stained with antibodies to both GFP and NDV proteins, most cells in the infected airway showed either GFP or NDV staining. To be certain that elimination of IFN-λ2 in our reporter mouse was not skewing our results, we compared total IFN-α and IFN-λ production in BAL samples from NDV-infected WT and reporter mice and saw no significant difference between the strains (Fig 5).

**Figure 5.**
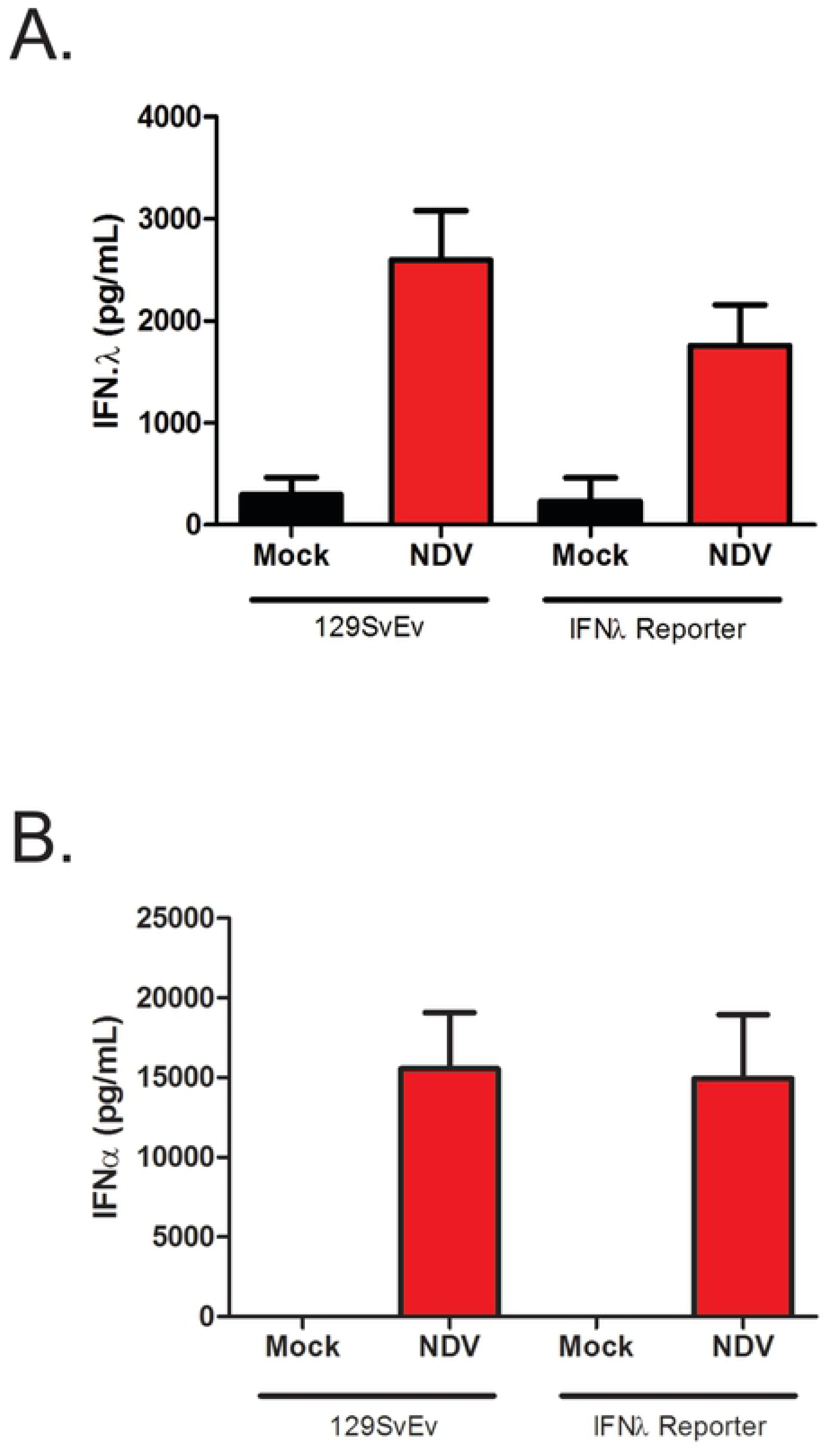
WT and IFN-λ reporter mice produce similar levels of type I and type III IFNs following NDV infection *in vivo*. (A) Levels of IFN-λ and (B) IFN-α proteins in BAL samples from WT or IFN-λ reporter mice 24 hours after mock- or NDV (10^7^ pfu) infection were quantified by ELISA. Data are pooled from two independent experiments.

### Immune cells are not a significant source of IFN-λ during NDV infection *in vivo*

IFN-λ synthesis by dendritic cells and monocytes exposed to viruses and pattern recognition receptor (PRR) agonists *in vitro* has been reported [7, 8, 11, 28]. So while results from our flow cytometry and histology studies using the IFN-λ reporter mice clearly demonstrated GFP expression from bronchial epithelial cells during NDV infection *in vivo*, we wished to determine whether there was also a contribution from innate immune cells in this model. To do this we used a panel of antibodies targeting cell surface markers to assess IFN-λ production, or GFP expression, by neutrophils (CD45^+^ F4/80^-^ Ly6G^+^), eosinophils (CD45^+^ F4/80^+^ SiglecF^+^ CD11c-), monocyte-derived macrophages (CD45^+^ F4/80^+^ SiglecF^-^ CD11c^+^) and alveolar macrophages (CD45^+^ F4/80^+^ Siglec F^+^ CD11c^+^) purified from the lungs of NDV-infected IFN-λ reporter mice 24 hours post-infection. GFP expression was minimal in all of these cell types (Fig 6), and therefore it is unlikely that they are contributing significantly to IFN-λ production in this setting.

**Figure 6.**
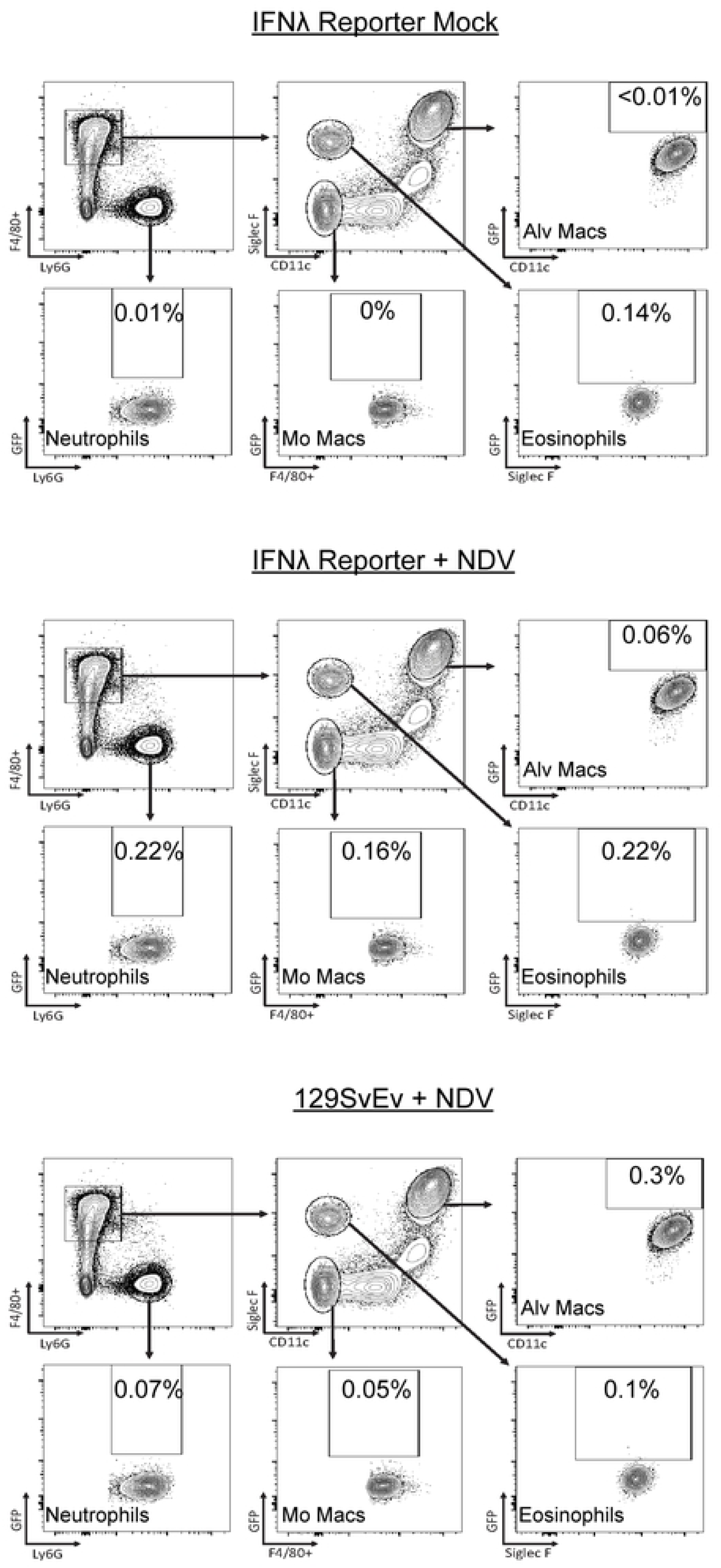
Alveolar macrophages and other myeloid cells do not produce IFN-λ during NDV infections *in vivo*. GFP expression was assayed by flow cytometry analysis of cells obtained from collagenase-digested lungs of mock-infected or NDV-infected IFN-λ reporter mice or infected WT animals. Cell populations are defined as follows: alveolar macrophages (F4/80+,Ly6G-, SiglecF+, CD11c+), neutrophils (F4/80-, LY6G+), monocyte-derived macrophages (F4/80+, Ly6G-, SiglecF-, CD11c-), and eosinophils (F4/80+, Ly6G-, SiglecF+, CD11c-). Data is representative of three independent experiments.

We were particularly intrigued by the lack of GFP expression in alveolar macrophages from NDV-infected IFN-λ reporter mice. Previously published reports using an IFN-α reporter mouse have identified alveolar macrophages as the major IFN-α-producing cell population during NDV and RSV infections *in vivo* [5, 6]. The results of our experiment demonstrate that alveolar macrophages do not contribute to IFN-λ production stimulated by NDV infection *in vivo*, suggesting that alveolar macrophages preferentially upregulate type I IFNs. To confirm this observation we carried out NDV infections in WT 129 SvEv mice and looked for induced IFN-λ expression in both epithelial cells and alveolar macrophages. Lungs from mock- or NDV-infected mice were harvested 24 hrs post-infection, and epithelial cells (CD45-EpCAM+) and alveolar macrophages (CD45+ F4/80+ SiglecF+ CD11c+) were sort purified by FACS. RNA from the sorted cell populations was used to synthesize cDNA for qPCR analysis which showed that IFN-λ transcripts were present only in the epithelial cell fraction isolated from NDV-infected mice (Fig 7). IFN-λ transcripts were not detected in the alveolar macrophage fraction from either mock-infected or NDV-infected mice, further confirming the readout from the IFN-λ reporter mouse strain. As an alternative approach to assessing IFN-λ production induced by virus, we used *in situ* hybridization to detect IFN-λ mRNA in lung sections from infected animals (Fig 8). IFN-λ RNA was found in bronchial epithelial cells of NDV-infected mice and not mock-infected animals. Taken together, the results from our qRT-PCR and *in situ* hybridization studies lead to the conclusion that only epithelial cells, and not alveolar macrophages, are responsible for IFN-λ production during NDV infection.

**Figure 7.**
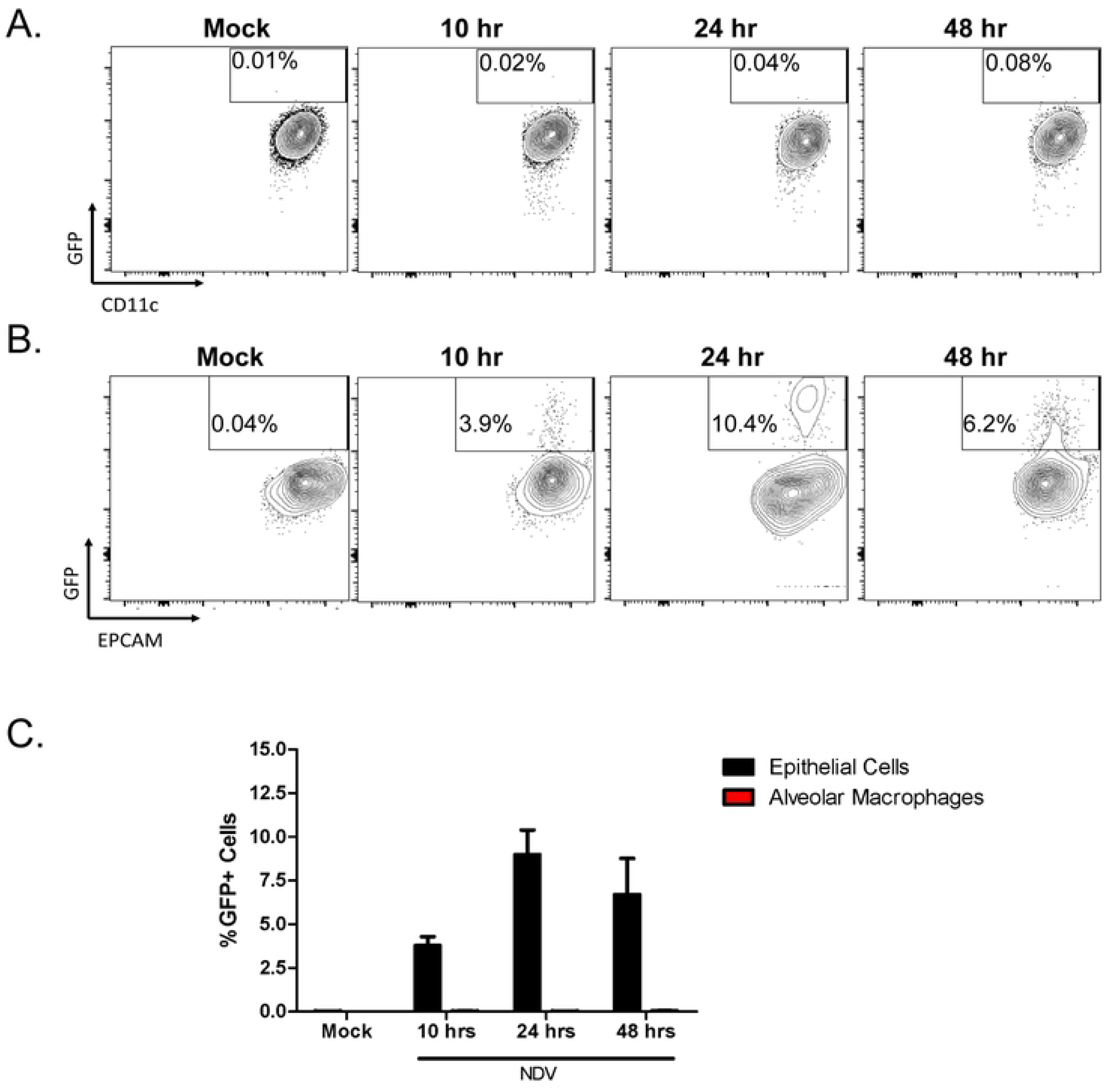
Kinetics of IFN-λ induction by NDV *in vivo*. GFP expression by (A) alveolar macrophages and (B) respiratory epithelial cells obtained from the lungs of mock- or NDV-infected IFN-λ reporter mice was assayed by flow cytometry. Samples were obtained at post-infection intervals of 10, 24, and 48 hours. (C) Graphical representation of mean % of GFP+ cells at each time point. Data are representative of two independent experiments.

**Figure 8.**
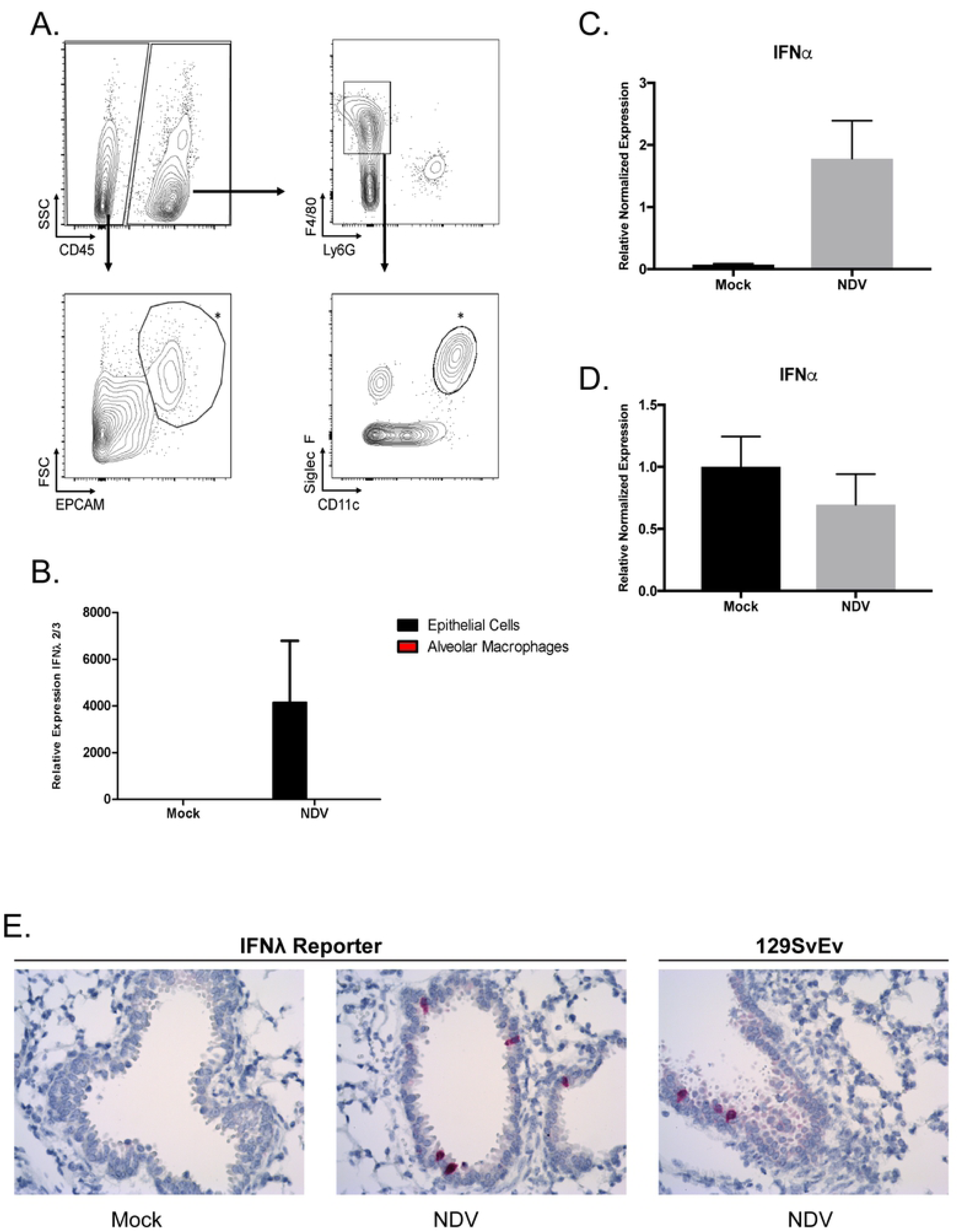
Distinct cell sources of IFN-α and IFN-λ *in vivo*. (A) Cell sorting strategy for isolation of alveolar macrophage and epithelial cell populations from collagenase-digested lungs obtained from mock- or NDV-infected WT mice 24 hours post-infection mice. Isolated populations of epithelial cells (CD45-/EPCAM+) and alveolar macrophages (as CD45+/F4/80+/Siglec F+/CD11c+) are indicated with an asterisk. (B) RNA harvested from FACS-sorted epithelial cells and alveolar macrophages and used to synthesize cDNA for detection of endogenous IFNλ2/3 transcripts by qPCR. qPCR detection of endogenous pan-IFNα transcripts (non-IFNα4) was also performed on FACS-sorted (C) alveolar macrophages and (D) epithelial cells. Expression is shown normalized to 18s rRNA. Data are pooled from two independent experiments. (E) *In situ* hybridization assays to detect endogenous IFN-λ transcripts were carried out on FFPE lung sections from NDV-infected WT and IFN-λ reporter mice at the 24 hour time point using probes specific for IFN-λ2/3 transcripts (seen in red). Images are representative of two independent experiments.

To ensure that our study produced results consistent with the work of Kumagai *et al*. [5], we also assayed RNA purified from the sorted epithelial cell and alveolar macrophage populations (Fig 8A) for the presence of IFN-α transcripts. As expected, induction of IFN-α mRNAs were observed only in alveolar macrophages (Fig 8C&D).

### IFN production by alveolar macrophages *ex vivo*

Noting the inability of the alveolar macrophage population to synthesize type III IFNs in the course of NDV infection, we wished to determine whether these cells were intrinsically unable to produce IFN-λ in response to NDV, or whether extrinsic factors in the lung microenvironment were restricting their production of IFN-λ. To ask this question, alveolar macrophages were purified from the lungs of naïve 129 SvEv mice by collagenase digestion and FACS sorting (Fig 9A) then mock-infected, or infected with NDV at an MO1=10, for 24 hours. The cells were then collected for RNA extraction and assayed by qPCR for the presence of IFN-λ transcripts. RNA isolated from a rodent cell line constitutively expressing FLAG-tagged mIFN-λ2, was used as a positive control. This analysis failed to detect IFN-λ expression from either mock-infected or NDV-infected alveolar macrophages (Fig 9B), a result confirmed by ELISA assay of media from the infected alveolar macrophage cultures (Fig 9C). Assay of the same samples showed high levels of IFN-α in media harvested from NDV-infected alveolar macrophages (Fig 9C). These results demonstrate that the same stimulus results in the production of either type I or type III IFN in a cell-type specific manner.

**Figure 9.**
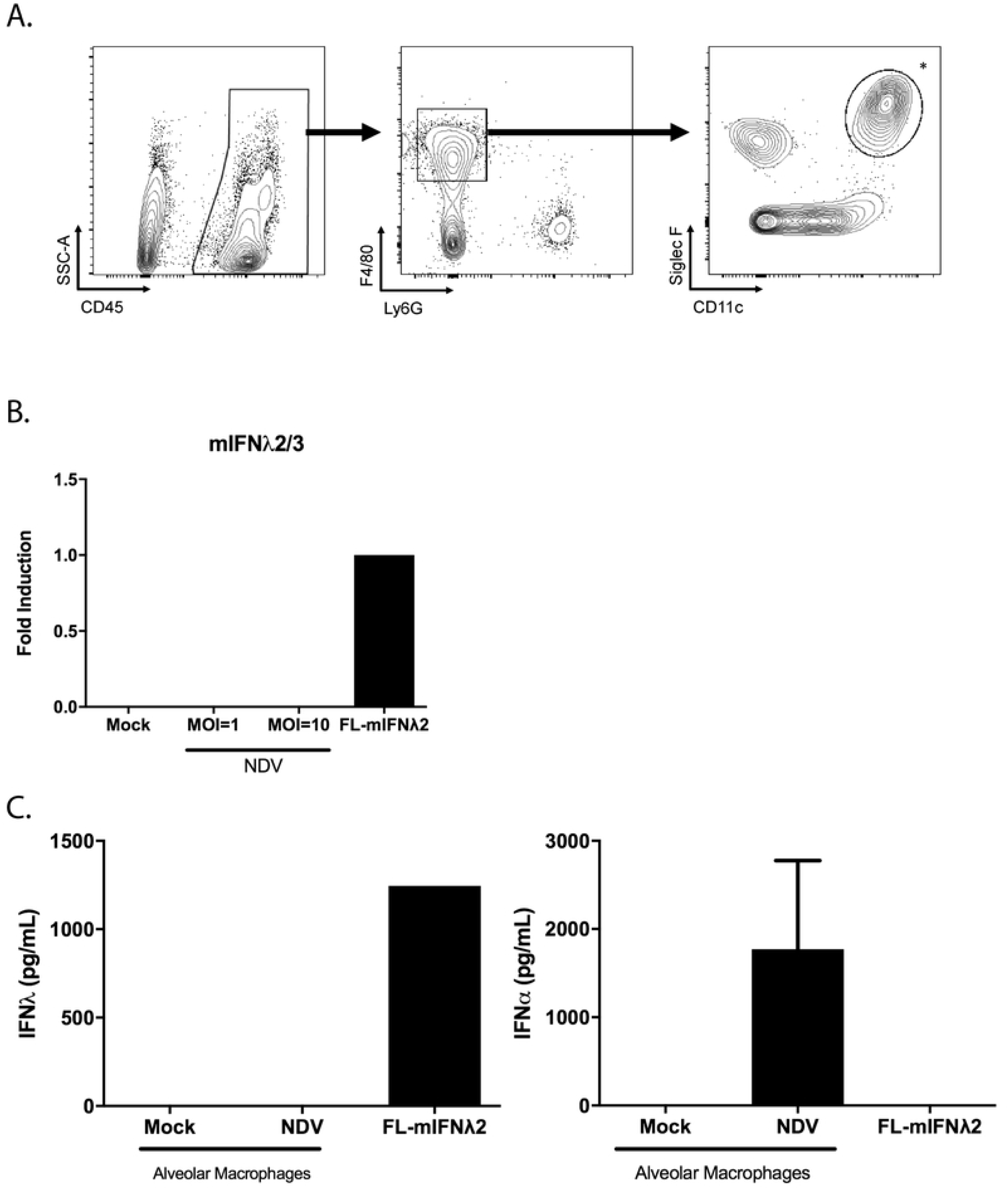
NDV infection of alveolar macrophages *in vitro*. (A) Sorting strategy for the isolation of alveolar macrophages from collagenase-digested lungs of uninfected WT mice. Isolated alveolar macrophages were then infected with NDV *in vitro* at the indicated MOIs. Supernatants and RNA samples were collected at 24 hours post-infection. (B) cDNA was synthesized from the RNA of mock- or NDV-infected cells and used to detect *Ifnl2/3* expression by qPCR. RNA isolated from cells stably transfected with a construct expressing FLAG-tagged murine IFN-λ2 (FL-mIFN- λ2 cells) was included as a positive control. (C) Levels of IFN-λ and IFN-α proteins secreted into the medium by cultured alveolar macrophages were determined by ELISA. Culture medium from FL-mIFN-λ2 cells was used as a positive control. Data are representative of two independent experiments

### Plasmacytoid dendritic cells, and not epithelial cells, are the major producers of IFN-λ during influenza A virus (IAV) infections *in vivo*

Intranasal administration of the non-replicating NDV allowed us to examine type I and type III IFN production following initial exposure, without regard to virus spread. However we also wished to use the *Ifnl2^gfp/gfp^* reporter mouse strain to characterize IFN-λ induction by influenza A virus (IAV), a virus capable of robust replication and spread in this species. Published studies have reported co-production of type I and type III IFNs from both pDCs [11] and epithelial cells [23, 29] exposed to IAV *ex vivo,* and we wished to characterize the *in vivo* source(s) of these cytokines. Lung homogenates from *Ifnl2^gfp/gfp^* mice infected with the WSN strain of IAV (10^6^ pfu) were prepared 48 hours post infection, and assayed by flow cytometry for GFP expression. By this method, measurable GFP expression was detected only in the pDC subset (CD45+/CD11c+/CD11b-/B220+/PDCA1+) and not in epithelial cells (Fig 10A-C).

**Figure 10.**
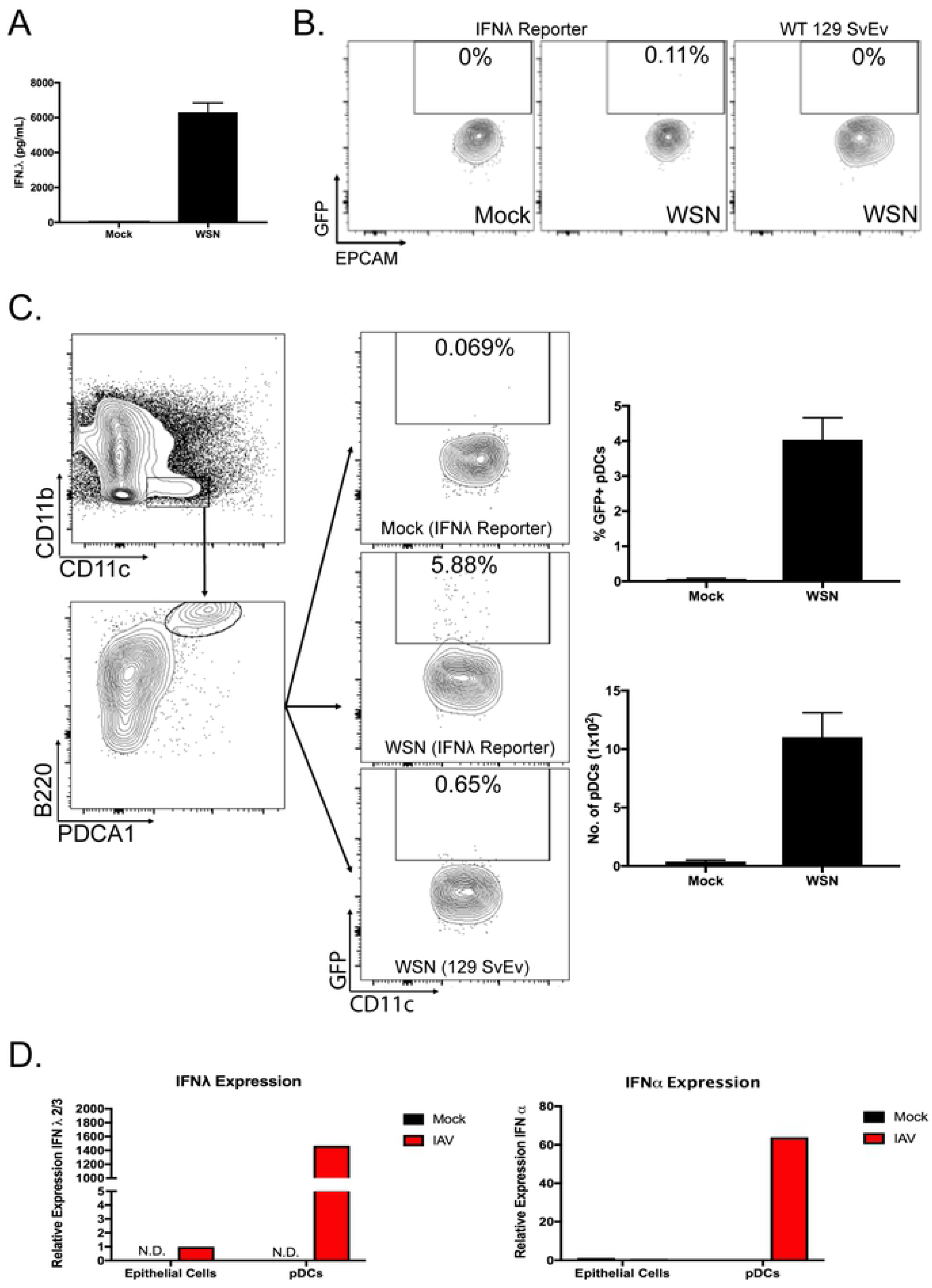
Plasmacytoid dendritic cells are the major producers of IFN-λ during IAV infections *in vivo*. (A) BAL samples collected from mock-infected or IAV-infected IFN-λ reporter mice were assayed by ELISA for the measurement of IFN-λ protein in airways. (B) Representative flow cytometry plots evaluating GFP expression specifically in epithelial cells, characterized as EPCAM+ cells (initially gated from DAPI-/CD45- and single cell populations), from single-cell suspensions of collagenase-digested lungs from mock-infected and IAV-infected (WSN; 10^6^ PFUs, 48 hrs) IFN-λ reporter mice or WT 129 SvEv controls. (C) Flow cytometry plots demonstrating gating scheme of pDCs from the lungs of the same mice as in (A). From initial gating of DAPI-/CD45+ and single cells, pDCs were gated as CD11c+/CD11b-, B220+/PDCA1+, from which GFP expression was assessed. Graphs on the right represent SEM of %GFP+ pDCs, and number of pDCs from each cohort. Data are pooled from three independent experiments. (D) qPCR analysis of IFN-λ and IFN-α transcripts in epithelial cells and pDCs that were FACS-sorted from the single cell suspension of collagenase-digested lungs from of mock-infected and IAV (WSN)-infected WT 129 SvEv mice (10^5^ PFUs, 48 hrs). Data are representative of two independent experiments. (E) Formalin fixed lung tissue sections from mock-infected or IAV-infected (10^6^ PFUs WSN) IFN-λ reporter mice, harvested at the indicated time points, were stained with IFN-λ oligonucleotide probes to detect *ifnl* RNA transcripts. Data are representative of two independent experiments.

Given the data obtained with NDV, and a recent study which concluded that epithelial cells are the primary source of IFN-λ in IAV infection [30], we considered the possibility that the more numerous epithelial cells might produce lower levels of IFN-λ on a per cell basis, below the level of detection by GFP expression, but still contribute to overall production of this cytokine. We approached this question in two ways. Using the sorting strategy shown in Figure 10 B & C, epithelial cells and pDCs from lungs of IAV-infected WT mice and controls were obtained, and RNA harvested from these populations was assayed by qRT-PCR for the presence of *Ifnl2/3* transcripts. As shown in Figure 10D, while IFN-λ2/3 mRNA could be detected in EpCAM+ cells from infected animals, pDCs appeared to be the predominant source of both IFN-α and IFN-λ. This result was consistent with ISH hybridization studies using formalin-fixed lung tissues form IAV-infected WT and reporter mice. Immunofluorescence studies showed diffuse positive staining for influenza antigen in airway lining epithelium, and oligonucleotide probes did detect rare IFN-λ2/3 expressing epithelial cells (Fig 10E) in animals infected with the WSN strain of IAV. A similar GFP expression pattern in pDCs and epithelial cells was observed by flow cytometry in animals infected with the PR8 strain, in the presence or absence of the NS1 gene (Fig 11) 48 hrs post-infection, but we were not able to detect *ifnl* transcripts by ISH in PR8-infected epithelial cells (data not shown).

**Figure 11.**
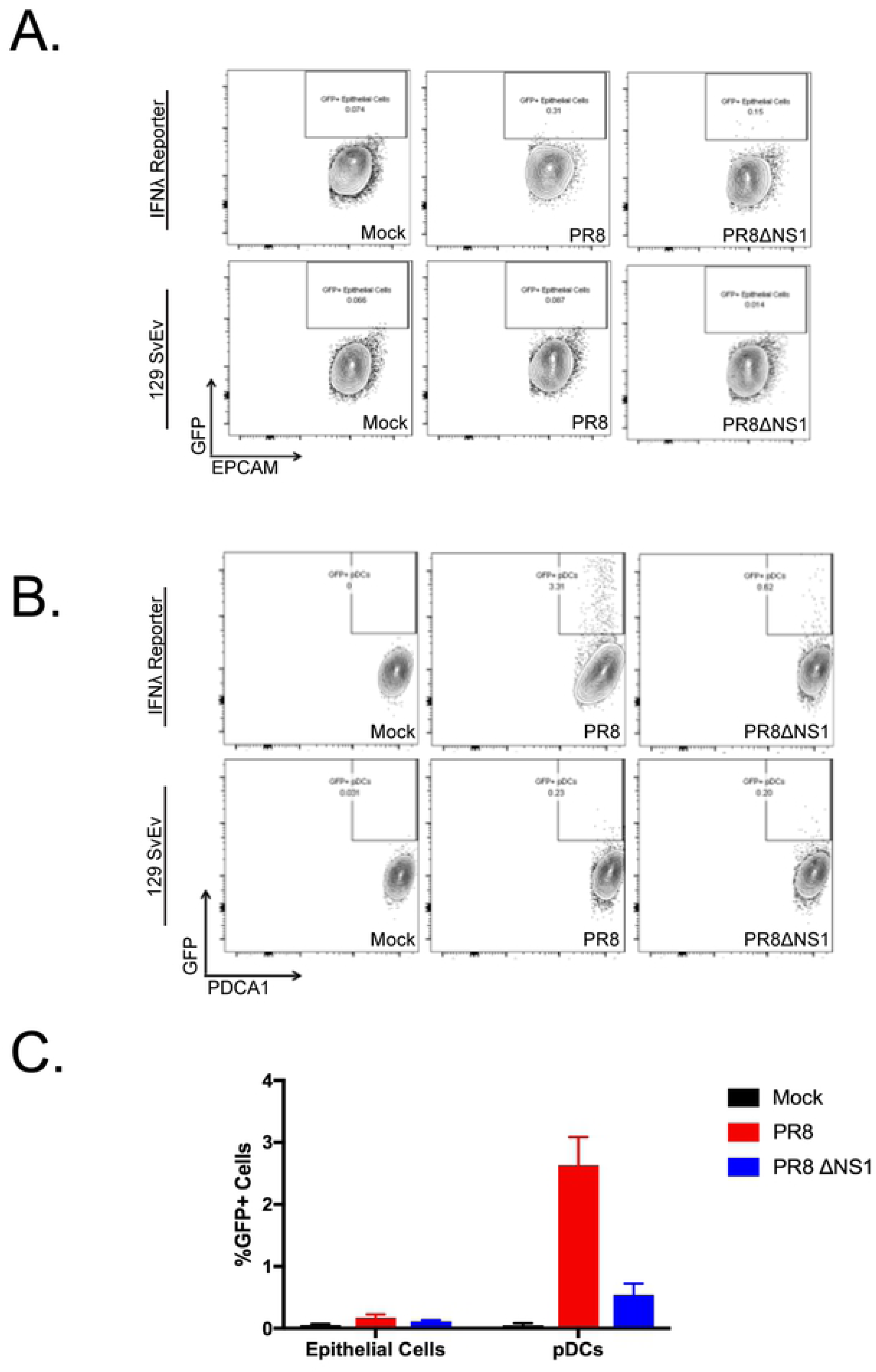
Minimal IFN-λ expression by epithelial cells during IAV infections is not due to a NS1-mediated inhibition. Representative flow cytometry plots showing GFP expression from (A) epithelial cells (gated from DAPI-/CD45-/EPCAM+) and (B) pDCs (gated from DAPI-/CD45+/CD11c+/CD11b-/B220+/PDCA1+) from wildtype 129 SvEv mice and IFN-λ reporter mice mock-infected or infected with 10^4^ PFUs of either IAV (PR8) or IAVΔNS1 (PR8ΔNS1) for 48 hrs. (C) SEM of %GFP epithelial cells and pDCs from each cohort.

To determine the relative contribution of these sources, we generated marrow chimeras using bone marrow harvested from newly generated IFN-λ ligand deficient mice, *ifnl-/-*, lacking both the *ifnl2* and *ifnl3* alleles (See Supplemental Fig. 2). Lethally irradiated WT mice were reconstituted with bone marrow harvested from either WT or *ifnl-/-* animals. Chimeric mice were then challenged with IAV, 10^5^ pfu of WSN, and BALs collected 48 hours post infection were assayed for the presence of IFN-λ by ELISA. IAV-infected mice reconstituted with *ifnl-/-* bone marrow showed significantly reduced levels of IFN-λ protein compared with IAV-infected animals which received WT bone marrow (Fig 12) demonstrating that ∼ 30%, of IFN-λ produced in response to IAV is derived from the epithelial compartment. To further pinpoint the source of this cytokine, WT mice were depleted of pDCs prior to IAV infection with anti-PDCA1 antibody, or a Rat IgG Isotype control (Fig 13A). Post infection measurements of BAL IFN-λ show a significant decrease in IFN-λ production in the absence of pDCs at 24 hrs post-infection, ∼ 60% (Fig 13B), consistent with the data obtained by generation of bone marrow chimeras.

**Figure 12.**
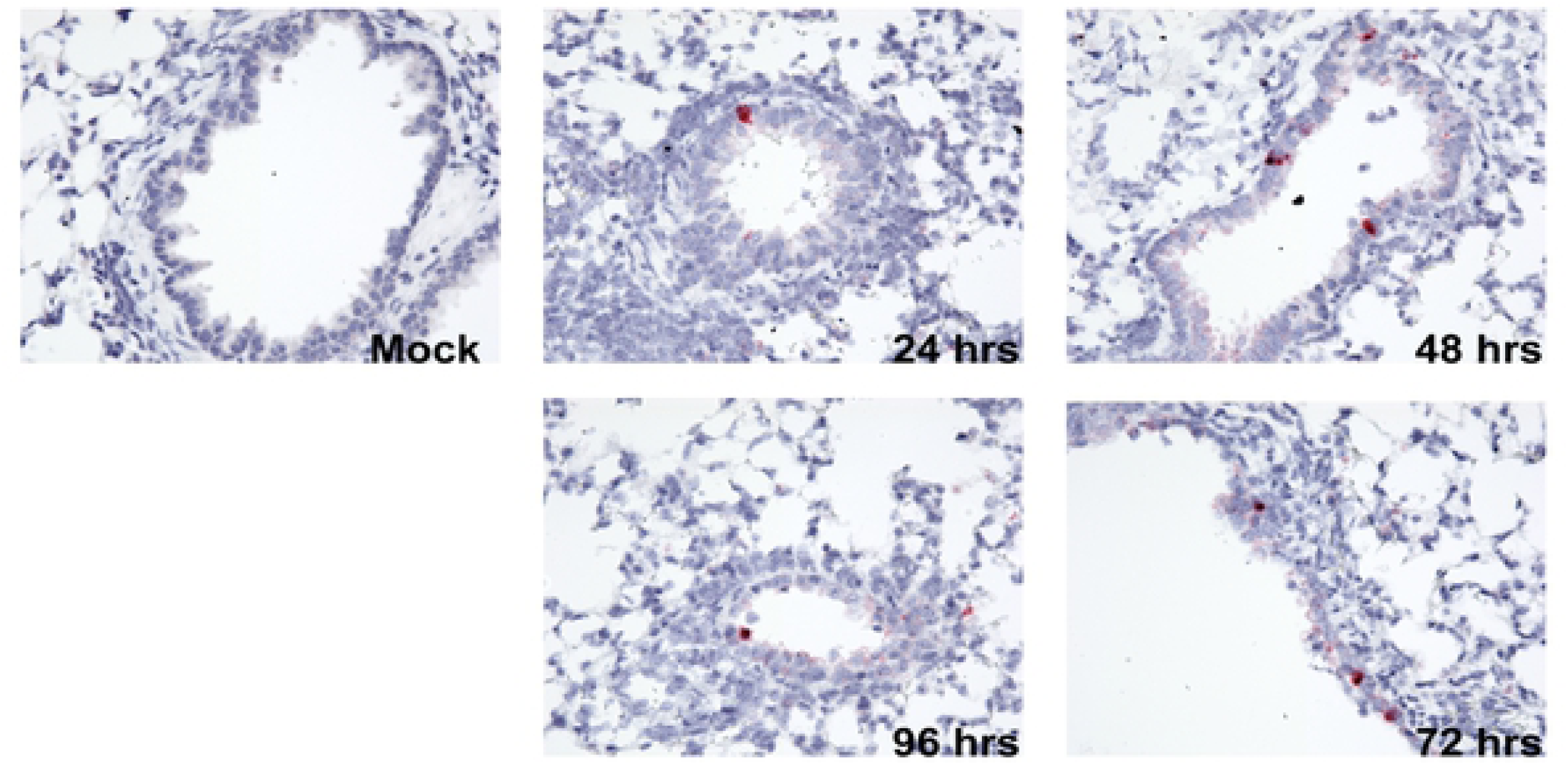
The hematopoietic compartment is the source of the bulk of IFN-λ protein produced during IAV infection. Bone marrow chimera mice, generated by reconstituting lethally irradiated wildtype C57BL6/J mice with bone marrow from either wildtype C57BL6/J mice or *ifnl*^-/-^ mice, were infected with 10^5^ PFUs IAV (WSN) for 48 hrs, at which point BAL samples were harvested and assayed for IFN-λ protein by ELISA. The graph represents the mean and SEM of IFN-λ protein levels measured from each cohort. Data were pooled from two independent experiments

**Figure 13.**
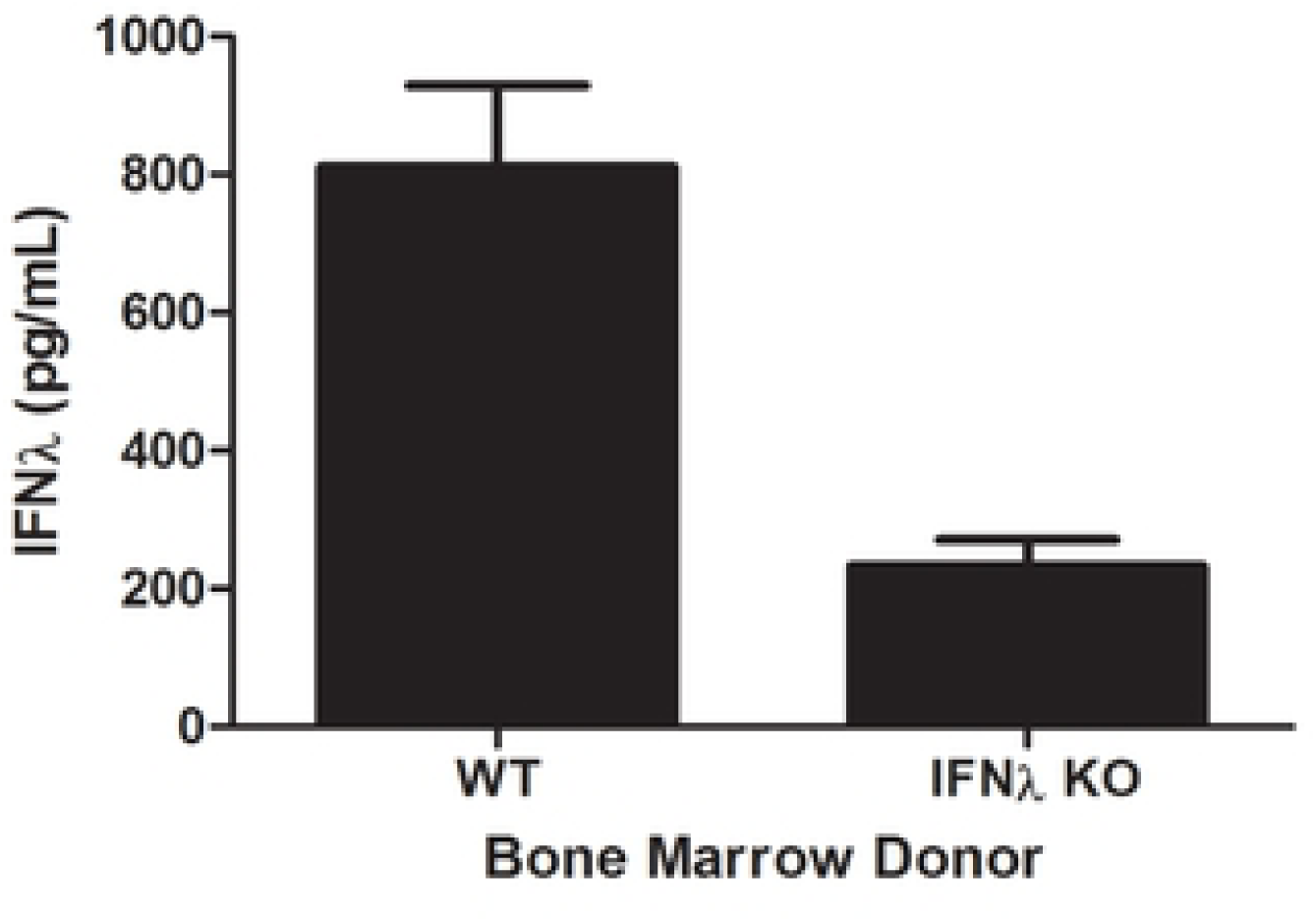
Depletion of pDCs during IAV infections in vivo results in significant reduction of both Type I and Type III IFNs. (A) Representative flow cytometry plots show pDC populations in the spleen and blood of IAV-infected mice (10^6^ PFUs WSN strain; 24 hrs post-infection) that were treated with the pDC-depleting antibody (anti-PDCA1) or the Rat IgG isotype 24 hours and 48 hours prior to IAV infection. Graphs on the right show mean percentages, and SEM, of pDCs in the spleen and blood of each indicated cohort. (B) BALs were harvested 24 hours post-IAV infection for IFN-λ protein measurements. Graphs represent data from two independent experiments

### Pathway activation by virus

As IFN-λ induction by NDV occurred only in CD45-EpCAM+ cells, and IFN-α induction only in alveolar macrophages in NDV infection, we hypothesized that production of either cytokine would depend on cytoplasmic PRRs. Many single stranded RNA viruses, including NDV and influenza [31], have been found to activate retinoic acid inducible gene-I (RIG-I). RIG-I bound to viral RNA interacts with the adaptor protein called mitochondrial antiviral-signaling protein (MAVS), triggering the events leading to IFN synthesis. To determine the role of MAVS activation in the production of each IFN type, we made use of MAVS-/- mice, available on the C57BL/6 background. WT and MAVS deficient animals were inoculated with NDV, and both IFN-α and IFN-λ levels were determined by ELISA of BALs collected 24 hours post-infection. As expected, there was no induction of either cytokine in the absence of MAVS, demonstrating that the events triggered by NDV recognition are dependent on cytoplasmic pattern recognition pathways in both cell types (Fig 14).

**Figure 14.**
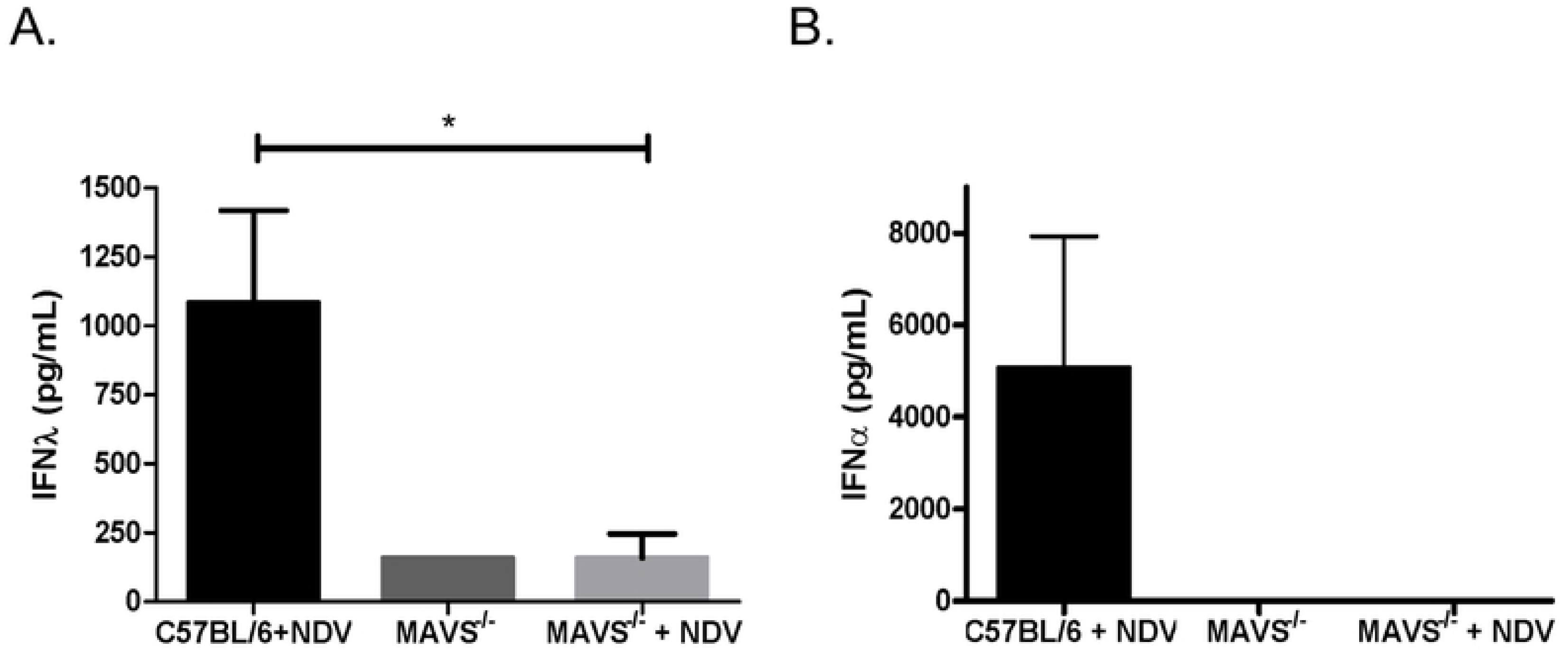
IFN-λ induction by NDV is MAVS-dependent. (A) IFN-λ and (B) IFN-α protein levels were measured by ELISA in BALs of mock-infected or NDV-infected (24 hrs, 10^7^ PFUs) WT and MAVS^-/-^ mice on the C57BL/6 strain background. Differences in IFN-λ levels between WT and MAVS^-/-^ mice were statistically significant as determined by student t test (P = 0.0474). No IFN-α protein was detectable in mock-infected or NDV-infected MAVS^-/-^ animals. Data were pooled from two independent experiments

Since our studies of IFN-λ induction by IAV suggested that pDCs are the predominant type III IFN producing cell population *in vivo,* we wished to determine which signaling pathways were triggered by this infection. Several reports have previously shown that pDC recognition of the influenza genomic ssRNA by the TLR7 endosomal pathway that signals through the protein myeloid differentiation primary response 88 protein (MyD88) [32–34] results in robust type I IFN expression. Based on our finding that pDCs are also the major source of type III IFNs in response to IAV infection, we suspected that this pathway was also required for optimal IFN-λ production. To test this assumption, MAVS-/- mice on the C57BL6 background were infected with the WSN and PR8 strains of IAV, and IFN-λ protein levels were measured in BALs. We saw no reduction in IFN-λ expression following PR8 infection of MAVS-/- mice at 24, 48 and 72 hours post infection (Fig 15A), but a ∼ 40% reduction following WSN infection at 48 hours post infection, consistent with data from our pDC depletion study. Conversely, in the absence of MyD88 (Fig 15B), a more significant reduction in IFN-λ was seen in the BALs of WSN-infected mice. Taken together, our data show that while both pathways can contribute to IFN-λ production following IAV infection, the major contribution is pDC derived, particularly for the PR8 strain of IAV.

**Figure 15.**
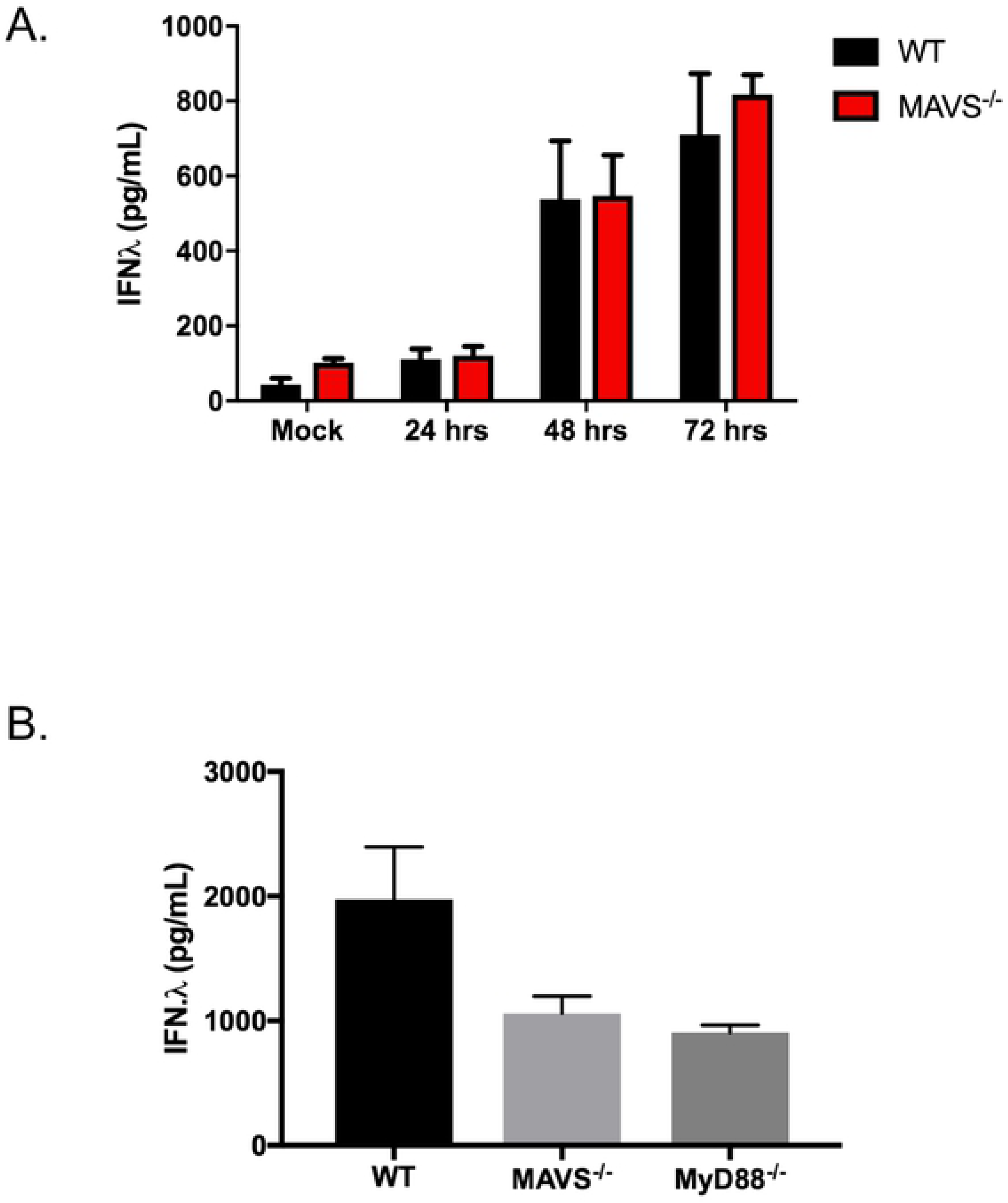
Contribution of MAVS and MyD88 signaling in IFN-λ induction during IAV infections is dependent on IAV strain. (A) IFN-λ protein levels were measured by ELISA from BALs of IAV-infected (10^6^ PFUs PR8) wildtype C57BL/6J mice or MAVS^-/-^ mice at the indicated time points. (B) IFN-λ protein levels measured by ELISA from BALs of IAV-infected (10^5^ PFUs WSN) wildtype C57BL/6J mice, MyD88^-/-^, or MAVS^-/-^ mice at 48 hours post-infection. Graphs represent the mean and SEM of IFN-λ protein measured from the BALs of each cohort. Data from each figure were pooled from two independent experiments.

We further explored this IAV strain dependence *ex vivo* using WT murine tracheal epithelial cells (mTEC) cultured at an air-liquid interface on transwell filters. IFN-*λ* protein levels in culture supernatants from virus infected mTECs showed robust production by NDV or WSN infected cultures, but no detectable IFN-*λ* following PR8 infection (Fig 16). This result is consistent with our inability to detect epithelial IFN-*λ* production following PR8 infection *in vivo*, and the absence of a requirement for functional MAVS protein.

**Figure 16.**
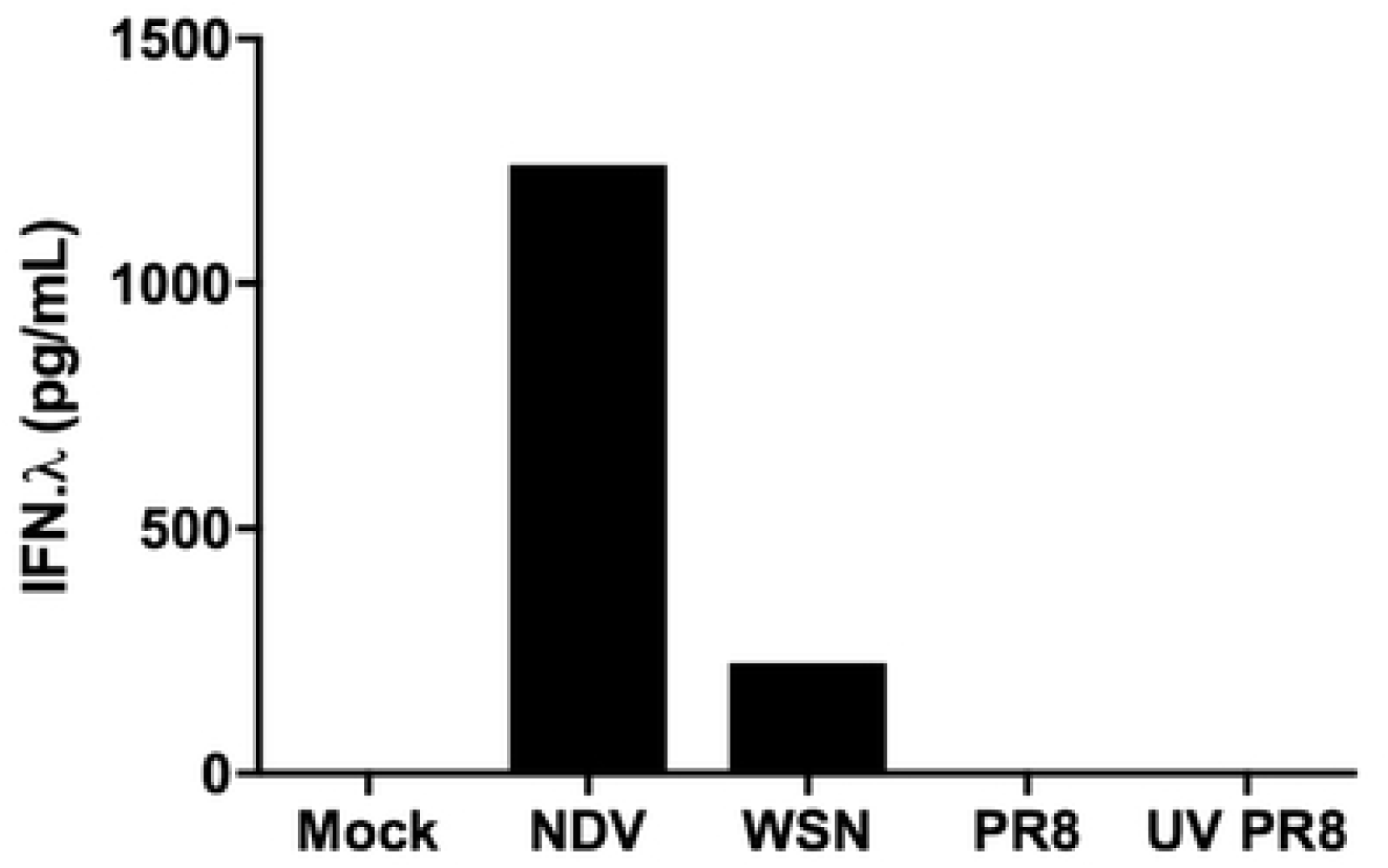
Murine tracheal epithelial cells produce IFN-λ in response to WSN but not to PR8 infection. Murine tracheal epithelial cultures were infected apically, at an MOI of 1, with NDV, WSN, PR8 or UV-inactivated PR8 for 24 hours. Supernatants were then assayed by ELISA for IFN-λ protein.

### Plasmacytoid DC production of IFN-λ requires type I IFN signaling

We and others have observed no significant difference in the resistance of wild type and IFNAR-/- mice to IAV infection [22, 35]. In the absence of the type I IFN pathway, IFN-λ production is sufficient to inhibit virus replication and spread. Assuming that pDCs were a major source of this cytokine, GFP expression was assayed in pDCs isolated from the lungs of PR8-infected *Ifnl2^gfp/gfp^* mice on WT or IFNAR-/- 129SvEv strain backgrounds. In this experiment, pDCs from IAV-infected *Ifnl2^gfp/gfp^* reporter mice with intact type I IFN signaling expressed GFP, but strongly reduced expression was detected in pDCs from IFNAR deficient *Ifnl2^gfp/gfp^* animals (Fig 17A). *In vitro* WSN infection of FLT3L cultured pDCs derived from bone marrow of wild type or IFNAR-/- mice produced the same result, with ELISA-detectable amounts of IFN-λ protein found only in medium from IAV-infected IFNAR^+/+^ pDC cultures.

**Figure 17.**
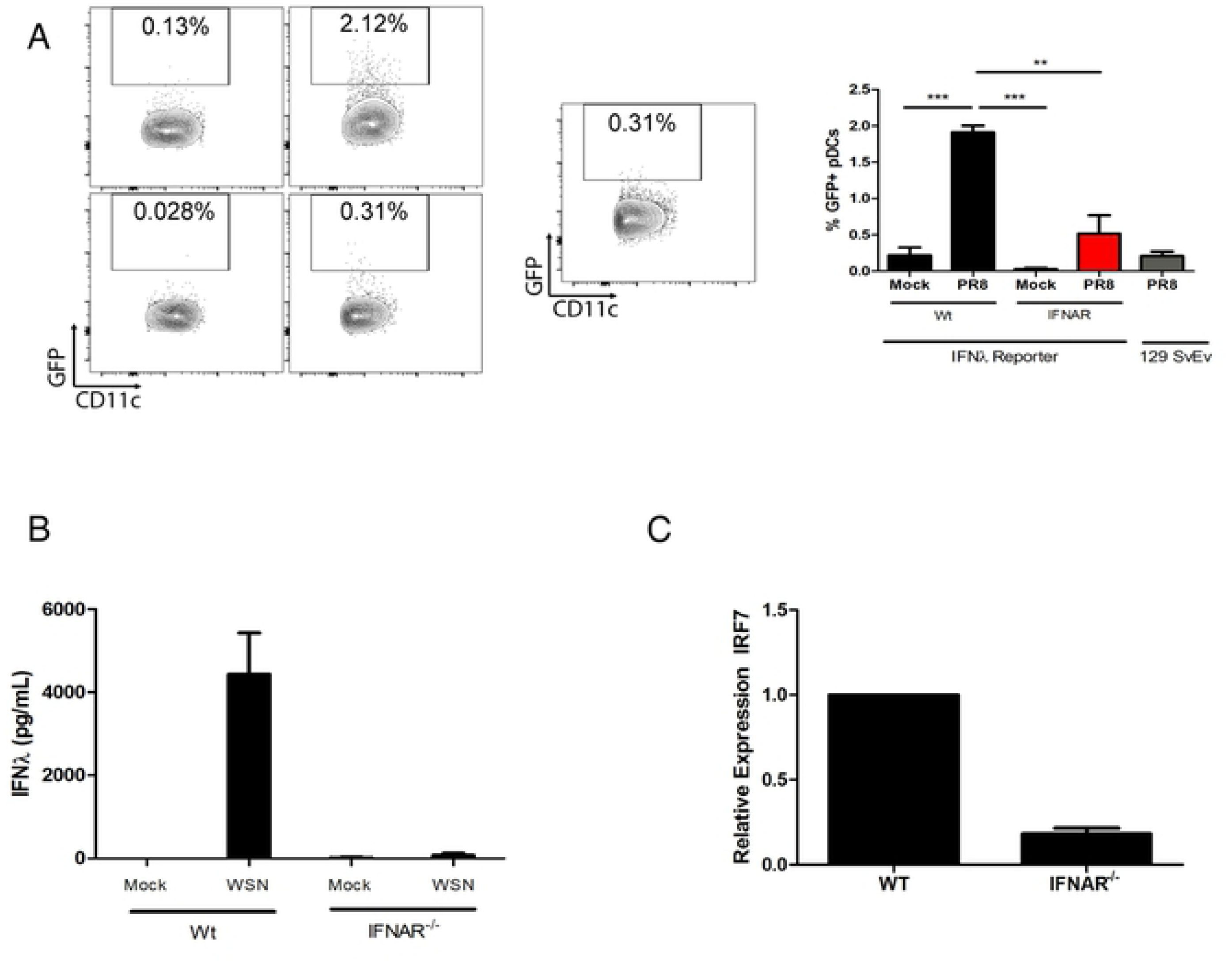
IFN-λ production by pDCs during IAV infection is dependent on type I IFN signaling. (A) Flow cytometry plots showing GFP expression in pDCs (DAPI-/CD45+/CD11c^+^/CD11b^-^/B220^+^/PDCA1^+^) from the lungs of IFNAR+/+ or IFNAR^-/-^ IFN-λ reporter mice, or 129 SvEv mice, mock-infected or IAV-infected (PR8 10^6^ PFUs, for 72 hrs). The graph on the right represents mean and SEM of %GFP+ cells from each cohort. (B) Supernatants from mock-infected or IAV-infected (WSN; MOI=1) WT and IFNAR^-/-^ FLT3L-cultured bone-marrow derived pDCs were collected 24 hours post-infection, and assayed for IFN-λ protein by ELISA. (**C)** qPCR measurements comparing basal levels of IRF7 mRNA expression between WT and IFNAR^-/-^ bone-marrow derived pDCs. Data were pooled from two independent experiments.

Plasmacytoid DCs are unique for their constitutive expression of interferon regulatory factor 7 (IRF7), a transcription factor essential for the induction of IFN-α genes [36]. IRF7 has also been implicated as a driver for IFN-λ transcriptional activation [37] [38]. Given the importance of IRF7 in the induction of IFN genes, we looked to see whether IRF7 levels were altered in the absence of IFNAR. As shown in Figure 17C, qPCR analysis demonstrated a substantial reduction in basal levels of IRF7 mRNA in IFNAR-/- pDCs. Since IRF7 is itself an ISG, we conclude that pDCs require some level of tonic activation of the type I IFN signaling pathway to maintain sufficient IRF7 for type III IFN induction in response to IAV infection. Interestingly, there was no decrease in GFP+ epithelial cells in NDV-infected IFNAR-/- IFN-λ reporter mice (Fig 18), supporting our conclusion that type I IFN-dependent maintenance of IRF7 levels in pDCs is essential for optimal type III IFN induction by IAV.

**Figure 18.**
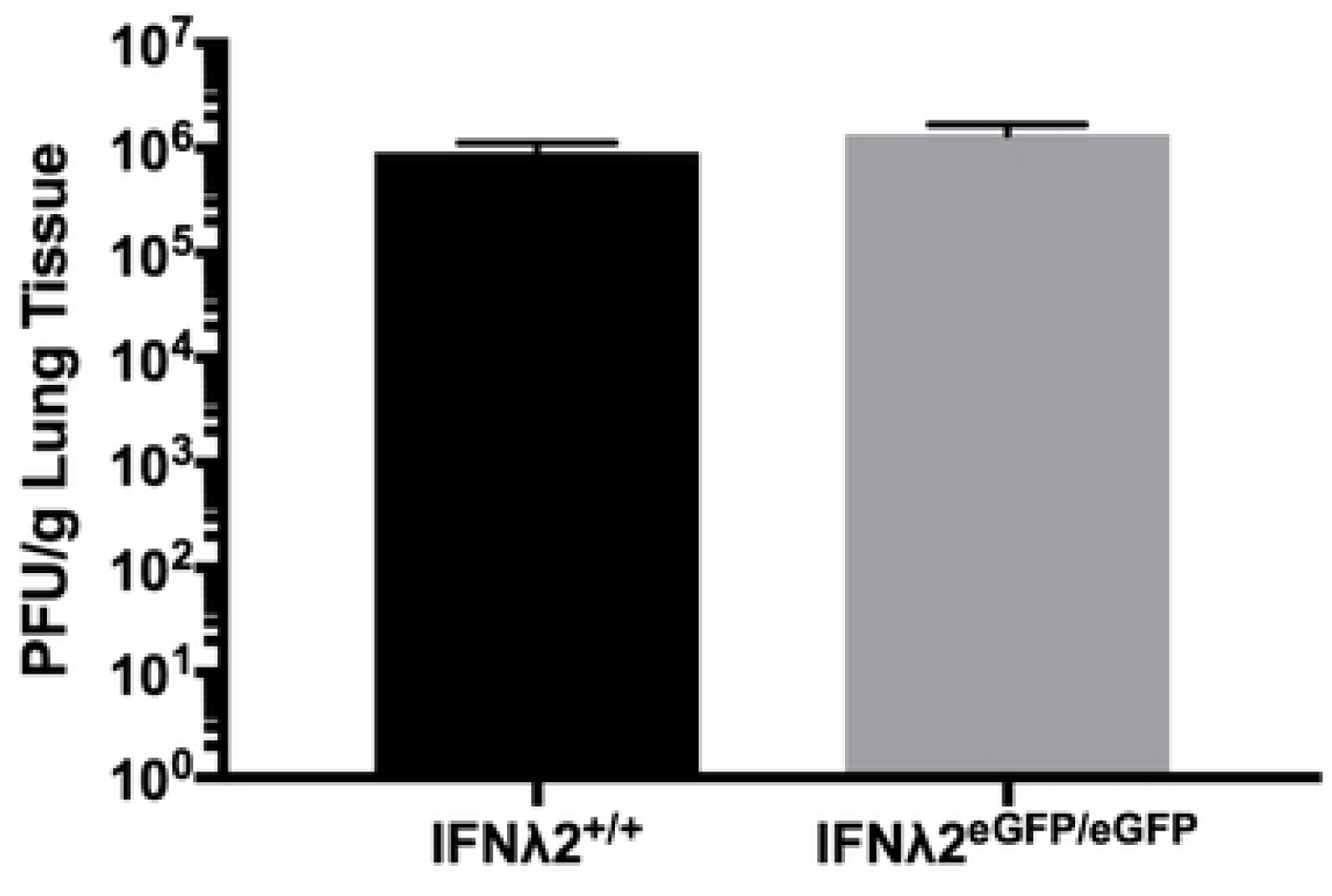
IFN-λ production during NDV infection is type I IFN-independent. GFP expression was assayed by flow cytometry of cells obtained from collagenase- digested lungs of 129 SvEv WT and IFNAR-/- IFN-λ reporter mice which were mock-infected, or infected intranasally with NDV for 24 hrs. Epithelial cells were defined as CD45-, EPCAM+. WT 129SvEV mice were included as controls for GFP expression, gating in the same manner. No significant differences in GFP expression were seen between IFN-λ reporter animals on the WT or IFNAR-/- backgrounds.

## Discussion

Since the discovery of type III IFNs in 2003 [3] [4] and the demonstration that identical sets of ISGs are induced by type I and type III IFNs in sensitive cells [39, 40], many groups have sought to determine whether these cytokines play a unique role in antiviral immunity. It has been found that type I and type III IFN receptors are differentially expressed by different cell types. While most immune cells in mouse and man are IFN-α responsive, only human pDCs, neutrophils and B cells have been found to consistently respond to IFN-λ [11, 30, 41, 42]. Until recently it was thought that all epithelial cells respond to both IFN-α/β and IFN-λ, but recent *in vivo* studies have demonstrated that intestinal epithelial cells become IFN-α insensitive soon after birth, and respond only to IFN-λ *in situ* [19]. These observations point to distinct roles for type I and type III IFNs in innate immune responses, determined primarily by the responsiveness and sensitivity of specific cell types to IFN-α/β and IFN-λ, but also by the availability of their ligands.

Our goal in generating an IFN-λ reporter mouse was to investigate the source of this cytokine as a viral infection of an epithelial surface progressed. This was done by replacing the *IFNL2* coding sequence with *eGFP*, a reporter gene whose expression is then regulated by the *IFNL2* promoter, UTR regions and other elements surrounding the *IFNL2* gene. Our first aim was to determine the extent to which GFP expression correlated with IFN-λ production by various cell types. *In vitro* virus infection of both epithelial and dendritic cells derived from the reporter mice induced GFP expression, and only GFP+ cells were found to express IFN-λ transcripts. Equally important were studies to confirm GFP and IFN-λ co-expression *in vivo*. Lauterbach *et al*. [17] had previously demonstrated that splenic CD8α DCs were the major producers of IFN-λ following systemic treatment with poly(I:C), a TLR3 ligand, and we established that GFP expression in the IFN-λ reporter mouse was limited to that cell type (Fig 3) following i.v. administration of poly(I:C). We next asked whether GFP expression by the *Ifnl2^gfp/gfp^* reporter mouse would recapitulate the pattern of IFN-λ expression previously described for heterologous rotavirus infection. Hernandez *et al*. [43] reported that IFN-λ transcripts were present only in CD45-EPCAM+ intestinal epithelial cells, but not expressed by cells isolated from lamina propria during RV infection. We repeated this study in WT and IFN-λ reporter suckling mice, harvesting tissue at the time point corresponding to maximal IFN responsiveness post-infection. Elevated, and equivalent, levels of IFN-λ were detected in intestinal homogenates from both strains following infection, with GFP expression detectable only in the epithelial compartment (Fig 6). We concluded from these results that GFP expression is an accurate indicator of IFN-λ expression in this reporter mouse strain and, importantly, that IFN-λ expression levels are equivalent in WT and *Ifnl2^gfp/gfp^* mice where production of IFN-λ3 protein compensates for the lack of the functional *IFNL2* gene.

The mucosal surfaces of the gastrointestinal and respiratory tracts are major portals of virus entry, and infection of these organs induces the secretion of both type I and type III IFNs [19, 22, 25, 43]. While both IFN types contribute to an effective antiviral response, receptor distribution suggests that type III IFNs have a primary role in protecting epithelial cells and that IFN-λ production is therefore crucial to an effective antiviral response at these barrier surfaces. While it is established that epithelial cells are the major IFN-λ producers during RNA virus infection of intestinal epithelial cells [10, 19], we wished to determine whether this was also a feature of respiratory virus infection. Using an IFN-α reporter mouse, Kamagai *et al.* [5] had demonstrated that IFN-α production following NDV infection was limited to alveolar macrophages. This was a surprising and important finding as it had been widely understood that essentially all cell types produced type I IFNs in response to NDV infection *in vitro*. As many respiratory virus infections result in the production of both type I IFNs and type III IFNs, we wished to determine whether alveolar macrophages were the source of both cytokines, or whether these cytokines were produced by different cell types in response to the same viral pathogen.

NDV infection of the IFN-λ reporter mouse resulted in GFP expression only by EPCAM+ epithelial cells as measured by flow cytometry and immunohistochemistry, and *in situ* hybridization studies detected IFN-λ transcripts only in bronchial epithelial cells of NDV-infected animals. This result is consistent with other published studies that have described IFN-λ production by mouse and human airway epithelial cells in response to respiratory viruses *in vivo* [23, 44–47]. Like Kumagai *et al*., we observed IFN-α mRNA induction in alveolar macrophages isolated from the lungs of NDV-infected animals, but no IFN-λ transcripts were detected in these cells. Therefore, NDV simultaneously induces both type I and type III IFNs upon respiratory infection, but each IFN type is produced by a different cell type.

Cellular recognition of RNA viruses generally occurs through recognition of viral genomes or viral transcripts by TLRs and RIG-I-like receptors, which signal through MyD88 and MAVS, respectively, to induce the expression of IFNs. TLRs are generally expressed by hematopoietic cells and RIG-I-like receptors by epithelial cells, although TLR3 expression at the apical surface of polarized murine and human tracheal epithelial cell cultures has been demonstrated [29]. Our results in MAVS-/- mice indicate that both IFN-α and IFN-λ production following NDV infection requires MAVS activation, but we do not yet understand how activation of this pathway by the same virus leads to production of one or the other IFN type in a cell type dependent manner. The most likely explanation is suggested by work showing that MAVS is located on the surface of mitochondria and peroxisomes [48], and further, that signaling via MAVS on peroxisomes induces type III, but not type I IFN production [9]. Preferential IFN-λ synthesis following MAVS activation was found to depend upon the relative abundance of peroxisomes in a given cell type, and polarization of intestinal epithelial cells resulted in increased numbers of these organelles. While these data predict a shift in the relative levels of type I and type III IFN production by different cell types, the complete absence of IFN-λ synthesis by alveolar macrophages in infected animals was unexpected. Further investigation is required to understand this compartmentalization of IFN production that we observe *in vivo*.

As NDV infection is abortive in mammalian cells [49], it was of interest to repeat these studies using IAV, which replicates and causes disease in the murine host. Based on our NDV data, and published reports which suggested that infected epithelial cells were the primary source of type III IFNs in IAV infection [30, 45], we expected our influenza studies to confirm this conclusion, but this was not the case. By all approaches taken it appeared that, for IAV infection, the bulk of IFN-*λ* was produced by pDCs rather than the respiratory epithelium. When pDC-depleted mice were infected with IAV, there was a 60% reduction in BAL levels of both type I and type III IFNs. Infection of bone marrow chimeras generated by the transfer of *Ifnl-/-* bone marrow into WT mice showed a similar decrease in type III IFN induction by IAV. In our study, ∼ 40% of the IFN-*λ* induced by IAV was from non-pDC sources, but this was dependent on virus strain. While GFP+ pDCs were detected using either the PR8 or WSN strains of IAV, only in WSN infections was epithelial IFN-*λ* synthesis detected *in vivo* or *ex vivo*. The conclusion that type III IFN synthesis induced by PR8 came primarily from pDCs was further supported by studies in MAVS-/- animals which showed no impact on IFN-*λ* levels in the absence of this pathway in PR8 infection.

Also of interest was our finding that IFN-*λ* production by IAV-exposed pDCs is type I IFN dependent. Our data suggest that the tonic signaling through the type I IFN receptor may be required to maintain the elevated basal levels of IRF7 which is a hallmark of this cell type [50]. While this result was unexpected, it is consistent with the observation that human subjects with a deficiency of IRF7 expression are more likely to experience life-threatening infections of influenza [51] as well as SARS-CoV-2 [52]. As both type I and type III IFNs are known to play a role in protection from respiratory viruses, this observation is further support of the hypothesis that pDCs are the major source of both of these cytokines during IAV infection. The demonstration that both IFN-*α* and IFN-*λ* are produced by IAV-exposed human pDCs [11] also supports this possibility.

In summary, we have used an IFN-*λ* reporter mouse model to determine the source of type III IFNs in two mouse models of respiratory virus infection. For NDV, type I and type III IFNs are simultaneously induced through the engagement of the same virus sensing pathway, but from two distinct cell types with respiratory epithelium as the sole source of IFN-*λ*. In IAV infection, pDCs appear to produce the bulk of type III IFN, and its production by this cell type requires type I IFN signaling. This study suggests that IFN induction by viruses *in vivo* is pathogen specific, as is the source of these cytokines.

## Methods

### Generation of the IFN-λ Reporter Mouse

Generation of the IFN-λ reporter mouse was carried out by ingenious Targeting Laboratory (www.genetargeting.com). The targeting vector was constructed using an 11 kb region of a C57BL/6 BAC clone (RP23: 24B20), containing the *Ifnl2* locus, sub-cloned into the pSP72 backbone vector (Promega). A GFP/FRT-flanked neomycin cassette was inserted into the BAC subclone using Red/ET recombineering technology, resulting in the complete replacement of the *IFNλ2* coding region of exons 1-5 (including intron sequences), with a long homology arm extending approximately 6.91 kb 5’ to the site of the cassette insertion and a short homology arm that extends approximately 2.68 kb 3’ to the site of the cassette insertion. The GFP-Neomycin targeting vector was linearized by Not I enzymatic digestion and electroporated into BA1 (C57B/6 x 129SvEv) hybrid embryonic stem cells. G418 resistant ES cell clones expanded for PCR analysis to identify homologous recombinants. Clones found to carry the GFP transgene were sequenced to confirm insertion, sequence fidelity, the genome/5’ cassette junction and the genome/3’ cassette junction. Southern blot analysis of positive clones was carried out using ApaI digestion of PCR products hybridized with a probe targeted against the 5’external region. Targeted BA1 hybrid embryonic stem cells were microinjected into C57BL/6 blastocysts and resulting chimeras with a high percentage agouti coat color were mated to C57BL/6 FLP mice to remove the neomycin cassette. Tail DNA from pups with agouti or black coat color was analyzed by PCR to confirm presence of transgene. Animals heterozygous for the *Ifnl2^+/gfp^* allele were backcrossed onto the 129SvEv background for 10 generations, then bred to produce homozygous animals. Genotyping was carried out by two PCR reactions: One reaction using Primer A (5’-CAGAGCTGGAAACTCAGAGCC-3’) and Primer B (5’-GACCGAGTCTGAGACCCACAAG-3’) and another reaction using Primer C (5’-CAGAGCTGGAAACTCAGAGCC-3’) and Primer B. Thermocycler conditions for those reactions were as follows: 95° C for 15 minutes, followed by 35 cycles of (94° C for 45 minutes, 65° C for 1 minute, and 72° C for 1.5 minutes), with an end step of 72° C for 5 minutes. The amplicon generated by Primers A and B, which encompasses the *Ifnl2* or *gfp* region, was then digested with the NcoI restriction endonuclease. NcoI treatment of the wildtype allele yields a 1100 bp band and a 700 bp band, while digestion of the amplicon containing the *Ifnl2^+/gfp^* allele, which lacks NcoI restriction sites, yields a single 1110 bp band.

### Additional Mice

Mice on the 129SvEv background lacking the type I IFN receptor (IFNAR-/-) were originally derived by Michel Aguet [53] and strain matched controls were purchased from Taconic Farms. MAVS-/- and MYD88-/- mice on the C57BL/6 background were obtained from Jackson Labs. Mx2-Luc Mx2 luciferase reporter mice were obtained from Hansjörg Hauser and Mario Köster [21]. Generation of the IFN-*λ*-/- mice, lacking both the IFN-*λ*2 and -*λ*3, sequences is described in supplemental figure 2. All mice were maintained under specific pathogen-free conditions in the vivarium of Rutgers-New Jersey Medical School.

### Cell culture

Bone-marrow derived FLT3-ligand dendritic cell cultures were generated from the bone marrow of IFN-λ reporter mice, *ifnl-/-* mice and WT controls. Bone marrow from femurs and tibia of 6-10 week old mice was washed, depleted of red cells with ACK lysis buffer, and cultured in RPMI+10% FBS containing 100 ng/mL human FLT3-ligand (Peprotech), 50 µM β-mercaptoethanol (Sigma), penicillin (100 IU/ml.) and streptomycin (100 μg/ml) at a cell density of 3×10^6^ cells/ml for 7 – 8 days.

For preparation of kidney epithelial cells, kidney capsules were removed, renal parenchyma was minced into 1 mm^3^ pieces and digested with collagenase IV (Worthington) at 37°C for 30 min. Following red blood cell lysis, digested tissue was pressed through a100 μM, then a 40 μM cell strainer, and the recovered cells were washed and then plated in DMEM/HamsF12 with 10% FBS. After 1-2 hours of incubation, non-adherent cells were collected and plated on collagen-coated dishes. Cells were allowed to reach 70% confluence prior to virus infection.

Murine tracheal epithelial cells were derived from WT or IFN-*λ* reporter mice at 6 to 10 weeks of age following the procedures outlined by You et al. [54].

### Poly I:C Treatment and Splenic DC Analysis

6-10 week old IFN-λ reporter mice were injected intravenously with sterile PBS or 100 µg polyI:C (InVivoGen). Splenocytes were harvested 6 hours after treatment, incubated with anti-mouse CD32/Fc-Block (BD Pharmingen) for 10 minutes at 4°C, then stained with the following antibodies: anti-mouse CD45-BUV395 (BD Horizon), anti-mouse CD3-APC Cy7 (BioLegend), anti-mouse CD11c-APC (eBioscience), anti-mouse CD11b-PerCP Cy5.5 (ebioscience), anti-mouse CD8α-V450 (BD Horizon), and anti-mouse Siglec-H-PE (eBioscience) antibodies. Analysis was carried out using the LSRII (BD Biosciences) flow cytometer.

### Virus infection and assay

Six-day-old suckling mice were infected with 4×10^6^ PFU of the simian RRV strain by oral gavage. Intestines were harvested 24 hours post-infection. These were either formalin fixed or homogenized in medium for virus quantitation. RRV quantitation was done using a focus forming assay as previously described [19].

Six-10 week old mice, under isoflurane anesthesia, were infected intranasally with 10^7^ PFU of NDV (Hitchner B1 strain) in 50 µl of PBS. Mice were euthanized 24 hours post-infection for collection of bronchoalveolar lavage fluid and lung tissue.

*In vitro* infection of primary kidney epithelial cells was carried out at a multiplicity of infection (MOI) of 2 and 20. Infections were done with RRV, the Hitchner B1 strain of NDV, the A2 strain of respiratory syncytial virus (RSV), and the WSN strain of influenza A virus (IAV).

A/PR?8?34 and A/WSN/33 influenza viruses were grown in embryonated chicken eggs. Allantoic fluid was assay for infectious viral particles by focus forming assay on MDCK cells. Serial dilutions of allantoic fluid were prepared and incubated on MDCK cells at 37°C, in 5% C0_2_ for 24 hours. Cells were then fixed in phosphate-buffered formalin, and viral plaques were visualized by incubating fixed cells with polyclonal rabbit anti-IAV serum (US Biologicals), followed by HRP-conjugated secondary antibodies for visualization of plaques via a colorimetric assay.

### Ethics statement

All mouse studies were approved by the Institutional Animal Care and Use Committees of Rutgers-New Jersey Medical School (protocols 999900760 and 13009C0316), performed in compliance with relevant institutional policies, local, state, and federal laws, and conducted following National Research Council Guide for the Care and Use of Laboratory Animals, Eighth Edition.

### *In vivo* luciferase detection

Six-day old Mx2-Luc mice were injected intraperitoneally with D-Luciferin (15 μg per pup), (Perkin Elmer), anesthetized with isoflurane, and imaged using the IVIS-200 Vivo Vision system (Caliper). Gray-scale images followed by bioluminescent images were then acquired and superimposed using the Living Image software (Caliper). Bioluminescent images represent luciferase signal intensity, with red as highest and blue as lowest, respectively. Regions of interest with detectable luciferase activity were measured quantitatively as relative light units (RLUs) at 3, 6, 12, 24, 48, 72, and 96 hours post-RRV infection.

### Immunostaining

Harvested tissues were fixed in 10% neutral buffered formalin, and submitted for processing and paraffin embedding. Deparaffinized sections underwent antigen retrieval and were subsequently blocked with Superblock (ScyTek) for 5 minutes at room temperature, washed in wash buffer (PBS+0.05% Tween) and then stained using anti-NDV antibodies (US Biological), anti-RRV antibodies (Meridian), anti-eGFP (Life Technologies) antibodies or anti-IAV antibodies (US Biological)overnight at 4°C. The following day, slides were washed in wash buffer and stained using anti-DyLight488 (Abcam) or anti-DyLight564 (Abcam) secondary antibodies for 1 hour at room temperature in the dark, washed in wash buffer, and incubated with DAPI for 6 minutes. After washing in distilled water, mounting medium (Vectashield) was applied, and slides were coverslipped. Images were obtained using an Axiovert 200M inverted fluorescence microscope (Zeiss) using the AxioVision LE64 software (Zeiss).

### Lung digestion and flow cytometry analysis

Perfused lungs were finely minced and digested in 5 mL of a 1.78mg/mL Collagenase IV (Worthington) solution in PBS (with Ca^2+^, Mg^2+^) for 45 minutes at 37°C. Lung digests were further dissociated by passage through a 20G needle, then filtration through a 100 µm filter. ACK lysis buffer was used to remove red blood cells, and remaining cell suspension was washed twice with PBS. Lung cell suspensions were first blocked with anti-CD32/Fc-Block (BD Pharmingen) for 10 minutes at 4°C, then stained with cocktail of primary antibodies including: anti-mouse CD45-BUV395 (BD Horizon), anti-mouse EPCAM-PECy7 (BioLegend), anti-mouse Ly6G-Alexa Fluor 700 (BioLegend), anti-mouse F4/80-APC (BioLegend), anti-mouse CD11c-PerCPCy5.5 (BioLegend), anti-mouse CD11b-APC-Cy7 (BioLegend), and anti-mouse Siglec F-PE (BD Pharmingen). Cells were then washed in FACS buffer (PBS+2% FBS and sodium azide), resuspended in FACS Buffer containing DAPI, and analyzed using an LSRII flow cytometer (BD Biosciences). Sorting of stained cell populations was carried out on an Aria II Cell sorter (BD Biosciences).

### Plasmacytoid DC depletion

Mice were injected i.p. with 500 μg of anti-PDCA1 or Rat IgG 24 and 48 hours before IAV infection. Mice were sacrificed 24 hours post-infection to collect BALS, and blood and spleens were collected to stain for pDC surface marker (CD11c+, CD11b-, B220+, PDCA1+).

### *In situ* hybridization

*In situ* detection of IFN-λ transcripts was carried out using the RNA Scope© HD 2.5 Detection Reagent-Red kit (Advanced Cell Technologies) according to manufacturer’s instructions. Briefly, formalin-fixed, paraffin-embedded tissue sections were heated at 60°C for 1 hour, deparaffinized in xylene and 100% alcohol, and incubated with hydrogen peroxide for 10 minutes at room temperature. Antigen retrieval was then carried out for 1 hour in RNA Scope antigen retrieval buffer, and slides were washed in distilled water and 100% ethanol, and then left to dry overnight. The following day, slides were washed in RNA Scope proprietary wash buffer and incubated in protease-plus buffer, washed again, and then incubated with mouse IFNλ2/3 DNA probes (Advanced Cell Technologies), followed by a series of RNA Scope adaptor probes, streptavidin-HRP, and chromogen solution to visualize IFN-λ transcripts. Slides were counterstained with 50% Lillie Mayer’s hematoxylin (American Master Tech Scientific Inc).

### RNA isolation and qRT-PCR

RNA from cells was obtained using the RNA Easy Kit (Qiagen) incorporating the DNAse I step to remove contaminating genomic DNA. cDNA was synthesized using iScript cDNA Synthesis kit (Biorad) using 10 ng RNA, with 10% of the cDNA product used for qPCR analysis. The following primer pairs were used to carry out qPCR reactions in a CFX96 Real Time System (BioRad) using SYBR Green (BioRad):

IFNλ2/3 Forward 5’-TCAAGC ACCTCTTCTCGATGG-3’
IFNλ2/3 Reverse 5’-AGCTGCAGGCCTTCAAAAAG-3’
IFNα Forward 5’-TCTGATGCAGCAGGTGGG-3’
IFNα Reverse 5’-AGGGCTCTCCAGACTTCTGCTCTG-3’
18s rRNA Forward 5’-GTAACCCGTTGAACCCCATT-3’
18s rRNA Reverse 5’-CCATCCAATCGGTAGTAGCG-3’
eGFP Forward 5’-GACGTAAACGGCCACAAGTT-3’
eGFP Reverse 5’-ATG CCGTTCTTCTGCTTGT-3’

Thermocycler conditions were set with the following parameters for amplification of IFNλ2/3 transcripts: 95°C for 3 minutes, followed by 40 cycles of 95°C for 10 seconds and a gene-specific annealing temperature for 30 seconds. Gene-specific annealing temperatures for IFNλ2/3, IFNα, eGFP were 58.4°C, 61.3°C, and 58.4°C, respectively. 18s rRNA qPCR reactions using the same annealing temperature as the target gene of interest were carried out simultaneously. Relative normalized expression was determined by normalizing amplification of IFN-λ, IFN-α, or eGFP expression to 18s rRNA levels by the ΔΔCt method [PMID 11846609].

### Data analysis

Data were analyzed using GraphPad Prism version 7 (graphPad Software). Mean and SEM were calculated by the same.

## Acknowledgments

Authors are thankful to Hansjörg Hauser and Mario Köster for Mx2-luciferase reporter mice. Histology and Imaging were done with the assistance of the New Jersey Medical Histology and Imaging Core Laboratory. This work was funded by grants from the National Institutes of Health: R21 AI073597 (JED), U01 AI082994 (JED) and R01 AI104264 (SVK and JED).

## Figure Legends

**Supplemental Figure 1.**
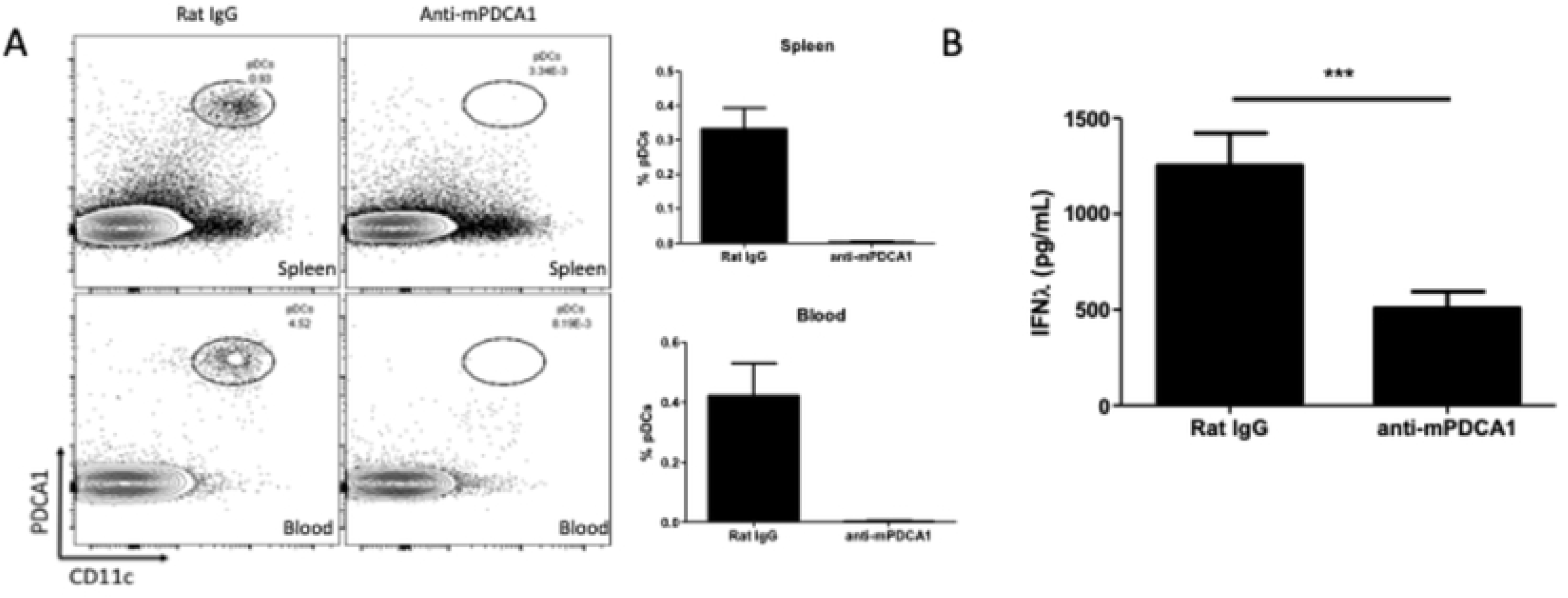
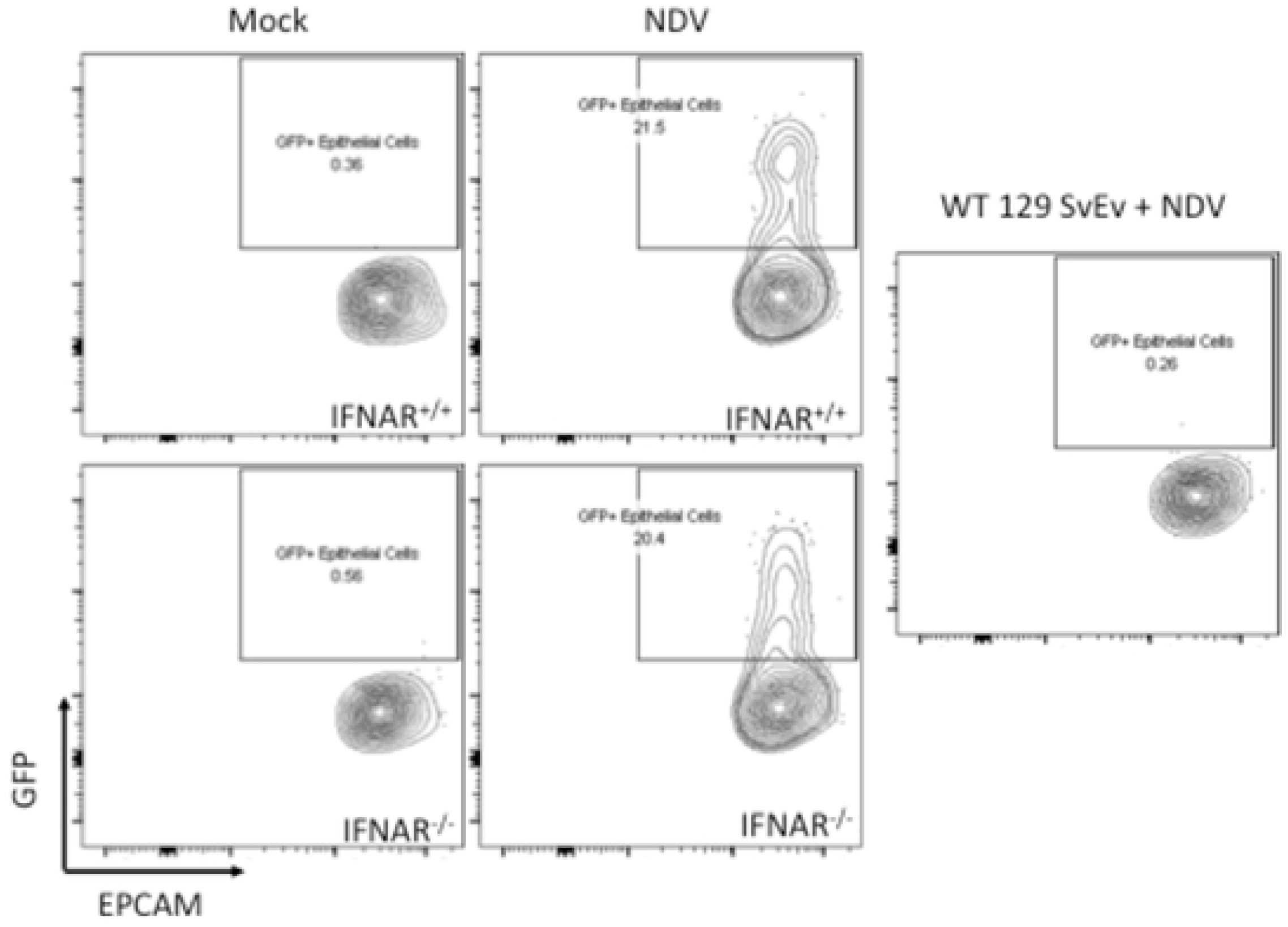
Viral loads are equivalent in WT and reporter mice. Cohorts of 6-10 week old WT 129 SvEv mice or strain-matched IFN-λ reporter mice were infected with 10^2^ PFUs of IAV (WSN). Viral titers from the lungs were assayed by immunofocus assay from lung homogenates collected at 72 hrs post-infection. The graph represents SEM of PFUs quantified for each cohort. This is experiment was performed twice.

**Supplemental Figure 2.**
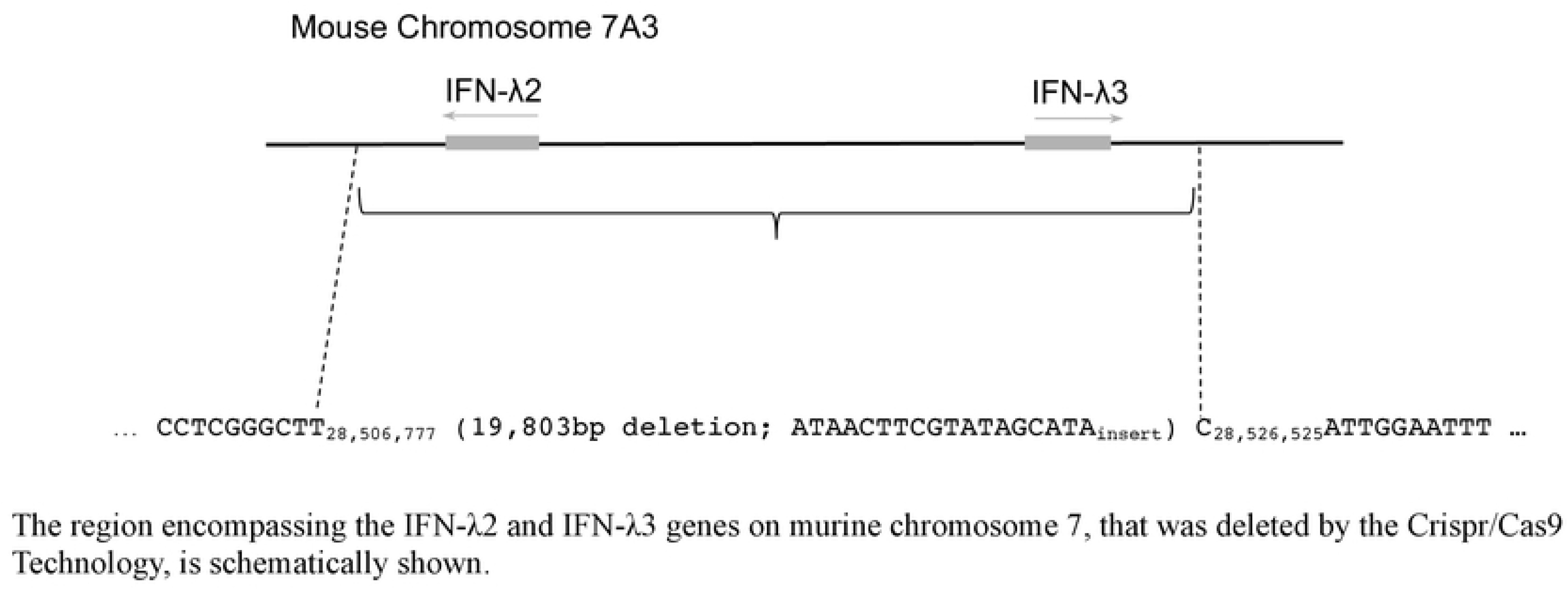
Generation of *ifnl-/-* mice. The mouse genome contains two functional IFN-λ genes, the IFN-λ2 and IFN-λ3 genes, which are juxtaposed in a head-to-head orientation on chromosome 7. Guidance RNAs (gRNAs) were designed to target sequences flanking these genes, and the CRISPR/Cas9 technology was used to generate mice lacking both IFN-λ genes, to create an IFN-λ2/3 knock-out (KO) strain. The selected IFN-λ2/3 KO strain contains 19,803 base pair deletion (from 28,506,772 to 28,526,525 nucleotide positions; NCBI GRCm38.p4) that removed the entire IFN-λ2 and IFN-λ3 genes including their promoters and 3’UTR, and replaced these with ATAACTTCGTATAGCATA sequence.

